# Dissolution of Phosphate and Precipitation of Carbonate in the Biomineralization of the Bivalve Shell Limnoperna fortunei

**DOI:** 10.1101/2024.05.30.596625

**Authors:** Antonio Valadão Cardoso, Rodrigo Novaes Ferreira

## Abstract

The mantle of bivalves plays a crucial role in the formation and maintenance of their shells through biomineralization. Detailed studies using scanning electron microscopy (SEM) and energy dispersive spectroscopy (EDS) analysis have revealed the presence of phosphorus (P) compounds as the primary phase during biomineralization at the growing edge of the periostracum of the bivalve shell Limnoperna fortunei (Dunker, 1857). The presence of a crystal morphology like hydroxyapatite (HAp) at the growing edge of the shell has also been identified, and the Ca/P ratio compatible with HAp. Carbonic anhydrase (CA), whose presence in the shell was investigated in this work, and/or bivalve proteins with identical capability are likely responsible for the dissolution phosphate and calcium carbonate precipitation. Other experimental techniques (ICP-OES, WDXRF) were used to quantify the main chemical elements in the shell of L. fortunei and the marine bivalve P. perna. The concentration of P in the shells suggests that phosphate is confined to the growing regions. FTIR and FTIR-ATR spectroscopies indicate aragonite as the main phase at the shell edges but also show the presence of phosphate absorption bands. X-ray diffraction (XRD) analyses revealed aragonite and calcite phases at the shell edges, with the presence of one of the main peaks of crystalline calcium phosphate both in L. fortunei and P. perna. The presence of phosphate as the primary phase in the biomineralization process of L. fortunei rekindles the discussion about the importance of the co-occurrence of phosphate and carbonate in the bivalve biomineralization dynamics and suggests an important evolutionary advantage in acquiring phosphate compounds essential for energy production and organism function.

## 1. INTRODUCTION

The control of golden mussel (Limnoperna fortunei, Dunker 1857) populations, which affect various economic activities in our country and in various parts of the world, has been the objective of different studies and research in Brazil (1). Funding Brazilian research has produced a set of fundamental studies indicating the incredible sophistication and complexity of the bivalve body. One example is the growth and biomineralization of the shell, which, to be understood, requires investigation of the living body that produces it, especially the mollusk mantle.

The shells of bivalves are composites produced through a complex biochemical system that constructs, in very short times, microscopic structures with mechanical properties that have deserved continuous attention (2–5). These shells have properties that, for centuries, have been admired for their beauty and mechanics. However, only with the advent of 20th-century analytical instruments, their submicroscopic structures could be partially understood (6). Several possibilities for advancing in obtaining new composites, inspired by biomineralization, have been presented in recent years aiming for superior mechanical properties associated with sustainability in the production process (7,8).

Bivalve biocomposites are currently produced in a matrix of calcium carbonate (CaCO_3_), predominantly in its thermodynamic phases of aragonite and calcite. Thirty years ago, Lowenstam and Weiner (1989) (9) suggested that the demand for phosphorus (P) and its compounds in biological processes would be among the main reasons why bivalves and different invertebrates biomineralize shells and other rigid parts in CaCO_3_ composites instead of calcium phosphate. Subsequent works supported this reasoning.

Phosphorus and its compounds are essential in the biochemical processes of living organisms. On Earth, all life is supported by P compounds, even viruses, as their membrane and genetic heritage are phosphorylated. DNA and RNA nucleic acids are phosphorylated; the main sources of energy reserves in organisms, including adenosine triphosphate (ATP), are phosphorylated as well. Many intermediate metabolites are phosphorylated compounds, and inorganic and organic phosphorus compounds are very suitable for biochemical syntheses and degradations. Several hypotheses have been raised about the biochemical option for P, including the ability of P compounds to be easily ionized at physiological pH(10).

Cook and Sheargold(11) suggested the occurrence of a massive phosphogenic event in the early Cambrian era which, associated with multiple environmental factors, resulted in a unique and remarkable phase in the evolution of life on the planet. It is considered that due to the availability of phosphates, shells during this period would mostly be made of calcium phosphate. A limiting factor for the growth of CaCO_3_ shells in phosphate-enriched oceans is the inhibitory effect that phosphate has on the nucleation and growth of amorphous or crystalline calcium carbonate phases by reacting with the substrate and blocking the carbonate growth(12). The presence of ionic phosphate would be inhibitory for carbonate growth at the shell tip.

As a limiting nutrient for primary productivity, phosphorus is rapidly recycled in the environment, limiting its accumulation in sediments(13,14) despite the difference in sedimentation velocity as compared to carbonate. Apatite, the most common phosphate mineral, has a molecular weight approximately 5 times higher than that of calcium carbonate. The higher molecular weight would favor sedimentation compared to calcium carbonate.

Thus, the advantage in producing phosphate shells was drastically reduced with the huge global expansion of the number of increasingly complex beings during the Cambrian period. Phosphorus is the eleventh most abundant element in the Earth’s crust, constituting approximately 0.1% of its weight(15). If the hypothesis of a more distant past consisting only of phosphate shells is correct, the inarticulate brachiopods can be considered “living fossils” because they are the only ones that currently build apatite-based shells containing also carbonate (16). Such composite shells are dramatically different from those based on CaCO_3_. Starting with the concentrations of calcium phosphate (approximately 50-60%), which are much lower compared to the concentrations of calcium carbonate commonly found in bivalve shells (approximately 95%). Recent work(17) with brachiopod shells indicates the existence of hierarchical structures, at the nano, micro, and macro levels, besides very sophisticated mechanical and rheological behavior (viscoelastic). These features are especially attributed to the concentration of non-biomineralized organic components. The presence of an amorphous organic phase in the shells seems to be essential for its flexibility and its ability to adapt to the narrow and irregular holes in which these beings live. The complex hierarchical organization of chitin+protein(17,18) organic fibrils and lamellae in the shells of inarticulate brachiopods may thus indicate evolutionary competencies.

By studying the compositional profile of brachiopod shells, Watabe and Pan in 1984 (19) observed that the concentration of calcium phosphates was higher near the periostracum. The subsequent layers consist of organic fibrils (chitin+proteins) with biomineralized veins of calcium phosphate particles. The innermost part would be entirely organic. They attribute the lower levels of phosphate in the innermost parts to the slow metabolism of these animals. To understand the source of shell phosphate, the same authors then studied the pathways of Ca and P absorption in the brachiopods Glottidia pyramidata (20) using radioactive isotopes. They concluded that the phosphorus in the shells originates from the food digested by the animal. But the work also presents an intriguing fact: a much higher concentration of ^32^P was found in the mantle rather than in the shell.

There are many reports on the presence of phosphate compounds in bivalve shells, but always in concentrations of ppm or ppb. Bevelander and Benzer (1948) (21) were among the pioneering workers who began to try to understand the calcification process and the formation of chitosan/protein/CaCO_3_ composites. They looked inside the mantle and were seeking to understand the internal dynamics until the materials reached its outer surface. In intentionally damaged shells, the authors observed that the first biomineralization granules were of calcium phosphate. Scholars, for over a century (W. Biedermann, 1914, cited by Bevelander and Benzer 1948), have already wondered why the mantle of modern bivalves contained calcium phosphate while the shell is built with calcium carbonate. Through the results obtained from the use of radioactive phosphorus (^32^P), Belevander in 1952 (22) suggested that the phosphorus present in the mantle is subepithelial and related to phosphorylation processes, whose function, the author found no connection with shell growth.

The presence of an extrapallial space proposed by A. de Waele in 1930 (23) between the mantle and the inner face of the shell made the discussion –about the phosphate present in the mantle even more suitable since the biomineralization activity would occur in that space therefore no longer directly from the pallial surface of the mantle (24). In the investigation of the components present in the extrapallial fluid, phosphate was not included (25). The attention given to the so-called extrapallial space as the site of chemical reactions resulting in the precipitation of CaCO_3_ seems to conflict with the nanometric dimensions (26) that this space would have, for example, in bivalves. In these animals, as far as we know, there is no experimental evidence to confirm whether this extrapallial space has a constant thickness or even if it is continuous. Moreover, the rheology of fluids, such as water and aqueous suspensions, confined between two hydrophilic surfaces of nanometric thickness, has specific physical characteristics (27). Thus, the so-called extrapallial fluid, which would fill this space, must have very different properties from a Newtonian fluid. Anyway, the scarce number of experimental measurements of the fluid and its cavity seems to produce more questions than answers (28,29).

Studies on mantle-shell interfaces in freshwater bivalves are notably scarce (29–31). Nonetheless, some authors suggest, relying on personal observations (29), that in freshwater mussels, the extrapallial space appears to be larger and more pronounced, entirely filled with extrapallial fluid. However, no conclusive evidence has been presented to support this claim.

The Limnoperna fortunei (golden mussel) is a freshwater bivalve and an exotic species in South America, where it arrived around the last decade of the last century(32,33). With very high reproduction rates (34), the adult golden mussel has millimeter-thick shells that, when dry, fracture and crumble with finger pressure. Observed under an electron microscope, the golden mussel shell presents a complex and hierarchical structure (35) with a first layer of calcite adhered to the periostracum, followed below by a nacreous layer of aragonite tablets, and the innermost part possessing a prismatic layer also of aragonite.

From the perspective of innovative new materials and bioengineering, the golden mussel, while causing economic losses by clogging pipelines, increasing maintenance costs for cooling systems, water intake screens, etc., also emerges as a kind of animal model for investigating the biomineralization process. This freshwater bivalve is a good candidate for investigating the structure of the mantle and the mantle-shell interface.

In the present work, we focus on the analysis of shell edges and their interaction with parts of the mantle. A long and thorough investigation using scanning electron microscopy (SEM) together with energy dispersive spectroscopy (EDS) technique on the complex structure of the mantle and its surface in contact with the L. fortunei shell was carried out to investigate the occurrence of calcium phosphate in the growing shell regions. Other experimental techniques, namely ICP-OES, WDXRF, XRD, FTIR, and FTIR-ATR were employed to confirm the presence of phosphorus (P) and phosphates at the growing shell edge. To verify if the occurrence of phosphorus and phosphate would be a singular case of L. fortunei, shell samples of P. perna were tested using the same techniques.

## 2. MATERIALS AND METHODS

### 2.1. Specimen collection and preparation

Specimens of Limnoperna fortunei (golden mussel) were collected from the reservoir of the Volta Grande hydroelectric plant, latitude 21°46’ South, longitude 42°32’ West, Minas Gerais, Brazil, and maintained in an aquarium in our laboratory.

Specimens of the marine mussel Perna perna (Linnaeus 1758) were acquired from the local market, originating from mussel farms in the region of Santa Catarina Island (located between latitudes 27°22′ and 27°50′ South and longitudes 48°25′ and 48°35′ West), in Santa Catarina, Brazil. In the laboratory, Perna perna specimens were kept frozen until used in future experiments.

For both species, shells were detached from the mussel body and were initially washed in a laboratory sink with running water until no traces of organic or inorganic materials (e.g., clays, sand, and other minerals) could be noticed. The shells were air-dried and stored in desiccator containers for future experiments. Sample preparations were carried out at the Laboratory of the Center for Bioengineering of Invasive Species in Hydroelectric Plants (CBEIH), located at Biominas, in Belo Horizonte. The preparation of the mantle-shell set for scanning electron microscopy (SEM) and energy dispersive spectroscopy (EDS) analyses were performed at the Microscopy Center of the Federal University of Minas Gerais, also in the city of Belo Horizonte, Minas Gerais, Brazil.

### 2.2. Quantitative chemical composition by ICP-OES

For the quantification of certain chemical elements present in the samples of L. fortunei (golden mussel) and Perna perna shells, the inductively coupled plasma optical emission spectrometry (ICP-OES) technique was used. This quantification was performed on three different occasions, in the year 2016, using an ICP-OES equipment, Perkin Elmer-Optima 3000 (Shelton, USA). In 2022 and 2023, we quantified again the metals present in shell samples using the Perkin Elmer DV8300 (Shelton, USA) equipment. The methodology used was validated according to the procedures recommended by official bodies (ISO-7 cleanroom and ISO-5 laminar flow hood). The tests were carried out at CIT-Senai in Belo Horizonte, MG.

### 2.3. Chemical composition by WDXRF

For the quantification of certain chemical elements present in samples of L. fortunei (golden mussel) and Perna perna shells, the wavelength dispersive X-ray fluorescence spectrometry (WDXRF) technique was used, Malvern Panalytical, model Axios Max (Malvern, UK). The tests were performed at CIT-Senai in Belo Horizonte, MG.

### 2.4. X-Ray Diffraction (XRD)

For the X-ray diffraction analysis of the crystalline phases present in the edges of the shells of L. fortunei and P. perna, a multi-purpose diffractometer, Malvern Empyrean Panalytical model (Malvern, UK) equipped with a copper tube and a two-dimensional PIXEL 2×2 detector was used. The experimental parameters of the measurement were 45 kV voltage, 40 mA cathode current, and a 2θ angle range of 20-90°. The quantification of the phases was performed based on the Inorganic Crystal Structure Database (ICSD). To perform the tests, the L. fortunei shells were carefully washed with deionized water and a brush, then air-dried.

After the cleaning step, the edges of the L. fortunei shells were easily broken manually with the aid of stainless-steel tweezers. Irregular pieces with millimeter sizes were obtained by this procedure. To collect the edges of the P. perna shells, it was necessary to use a steel clamp due to higher resistance to fracture of the shells. The tests were performed on the shell powder, ground with a mortar and pestle.

### 2.5. Scanning Electron Microscopy (SEM) with Energy Dispersive Spectroscopy (EDS)

Preparations and analyses of L. fortunei shells using scanning electron microscopy (SEM) were conducted at the Federal University of Minas Gerais Microscopy Center (CM-UFMG) using two instruments: a– FEI Quanta 3D FEG dual-beam (Hillsboro, USA) with detectors for backscattered electrons (BSE) and energy-dispersive X-rays (EDS); and b– APREO-2C ThermoFisher (Waltham, USA) with standard BSE and EDS detectors.

Ten specimens of L fortunei were selected, treated with buffered Karnovsky fixative solution, dehydrated with crescent alcohol concentrations and dried using the critical point drying Leica, model EM CPD030 (Wetzlar, Germany). Two types of preparations were investigated: a– Shells with all internal parts of the animal; b– Shells only with adhered mantle. For the preparation of the mantle-shell assemble, the specimens were opened with the help of a stainless-steel spatula and the body parts of the bivalve were delicately sectioned to ensure that the mantle-shell set could be clearly analyzed. After this initial stage, the samples were coated with 50 nm of graphite and kept in a low humidity desiccant cabinet prior and after the sessions.

Fourteen sessions of 2 hours each were carried out to investigate different parts of the L. fortunei shell and mantle. The maps of the chemical elements carbon (C), oxygen (O), calcium (Ca) and phosphorus (P) show slightly different colors in different figures due to SEM-EDS sessions carried out on different dates.

### 2.6. FTIR and FTIR-ATR Spectroscopy

FTIR and FTIR-ATR tests were performed on samples of the edges of L. fortunei and P. perna shells with identical protocols. To perform the tests, both L. fortunei and P. perna shells were carefully washed with deionized water and a brush, then air-dried. After this cleaning procedure, the edges of the L. fortunei shells were broken manually with the aid of stainless-steel forceps. Irregular pieces with millimeter sizes were obtained by this procedure. To collect the edges of the P. perna shells, it was necessary to use a steel clamp due to the higher resistance to fracture of the shells. Irregular pieces of these edges were pulverized for the FTIR tests. For the FTIR-ATR tests, entire irregular pieces of the edges of the shells were used.

Absorption infrared spectra were measured on a Perkin-Elmer FTIR model RX I spectrometer (Shelton, USA) between the wave number region 400–4500 cm^−1^. The powder samples were mixed with KBr and then pressed into a pallet. All measurements were carried out at room temperature. Powder spectra were recorded with a resolution of 2 cm^-1^ while the FTIR-ATR spectra, with a resolution of 1 cm^-1^.

### 2.7 Carbonic Anhydrase Assay

The presence of the enzyme carbonic anhydrase in L. fortunei shells was assessed using the protocols, with modifications, of Miyamoto(36) (method 1 from now and onwards) and Fuchs(37) (method 2), following the Sakalauskaite proposed preparations and cleanings (38). The method involves changing the pH from 8.3(method 1)/8.2(method 2) to 7.3(method 1)/6.5(method 2) due to the reaction of the metalloenzyme that catalyzes the interconversion of CO₂ and water into a bicarbonate ion and a proton. This method was developed by Freeman and Wilbur(39) and detailed by Wilbur and Anderson(40). 40 grams of L. fortunei shells were ground and left for 24 hours in a 2% sodium hypochlorite solution for cleaning. After air-drying, 20g were immersed in a 0.5M EDTA pH 8 solution (PHT, Belo Horizonte, Brazil) and 20g in 4% acetic acid (Vinagre Dicasa, Contagem, Brazil) for 6 days for demineralization, under continuous stirring. The supernatants were collected, diluted with distilled water, and refrigerated at 2°C for 24 hours, then centrifuged at 12,000 rpm for 20 minutes and stored again under refrigeration.

Half of the supernatant demineralized with EDTA received 2 mM ZnCl₂ to neutralize the metal chelating effect, followed by centrifugation and refrigerated storage. The other half was not. Bovine carbonic anhydrase (CAB, Sigma-Aldrich) at 12 mg/L in distilled water cooled to 2°C was tested as a reference. The amount injected in the tests was 1 ml of solution.

Following protocol **method 1** with modifications, 10 ml of pH 7 phosphate buffer (Dinâmica, São Paulo, Brazil) and 5 ml of the demineralized suspension and 6 drops of phenol red (Dinâmica, São Paulo, Brazil) were added to a 100 ml glass beaker, that placed in a Styrofoam container with cold water and ice. The addition of pH 10 buffer (Dinâmica, São Paulo, Brazil) raised the pH to 8.3. A digital pH meter (Akso, S Leopoldo, Brazil) with temperature control was inserted into the beaker and then, starting the test, 5ml of ice-cold water (0-2°C), previously bubbled with pure CO₂ for 30 minutes via a flowmeter (Voha Instr., Changzhou, China), was added to the beaker.

Using the **method 2** (37) with modifications, 40 ml of 0.1 M Tris buffer solution (Orion, São Paulo, Brazil), adjusted to pH 8.2 with the addition of H_2_SO_4_, were placed in a 100 ml glass beaker with a digital pH meter (Akso) and a stainless-steel CO₂ diffuser (commercialized by BoB, São José dos Campos, Brazil). Then the solutions investigated were added (5 ml of the demineralized suspensions or 1 ml of CAB), without exceeding 40 ml of liquid. The test was carried out at room temperature (23-25°C, Belo Horizonte, Brazil) and started with the injection of CO_2_ into the beaker using a flowmeter (Voha Instr., Changzhou, China.) set at 400 ml/min of CO₂.

## 3. RESULTS

### 3.1. SEM-EDS of the shell edges

The SEM images in Figure 1 were composed to show the entire area of the two shells with parts of the mantle of the bivalve L fortunei. SEM images of the two shells with the viscera of the golden mussel are presented in supplementary material S2. From the images in Fig. 1 it is possible to locate and visualize inside the mantle (see also S3 in the supplementary material) evidenced by the tears in the face. The inner part of the mantle has a 3D structure of interconnected fibers or aggregates of fibers, forming a micrometric “sponge-like” structure, which has already been presented in previous work on the mantle of other bivalves (41). The shells are approximately 2 to 3 cm long. Regions A1 are marked on the edges of the shell in Fig. 1.a, and A2, A3 and A4 are marked in Fig. 1.b. These areas were studied in detail to take images, collect maps of the chemical elements and perform semi-quantitative analysis using EDS spectroscopy. The A4 region is one of the places where tears in the face of the mantle make it possible to investigate its interior. Just above A4, structures called mantle edges can be seen on both shells (42) [43]. Of the three folds at the edges of the mantle (inner, middle, and outer) we have located the inner fold, which is visible, outlined and marked with the letter B in Figures 1.a and 1.b. The periostracum is excreted in the outer fold, its beginning can be in the image as a ledge adjacent and parallel to the inner fold. The inner face of the mantle is intact in some areas, torn in others. In the frayed areas, it is possible to see inside, including the inner side of the mantle’s outer face, which is in contact with the shell. In some areas it is the biomineralized surface of the shell itself that appears.

**Figure 1.**
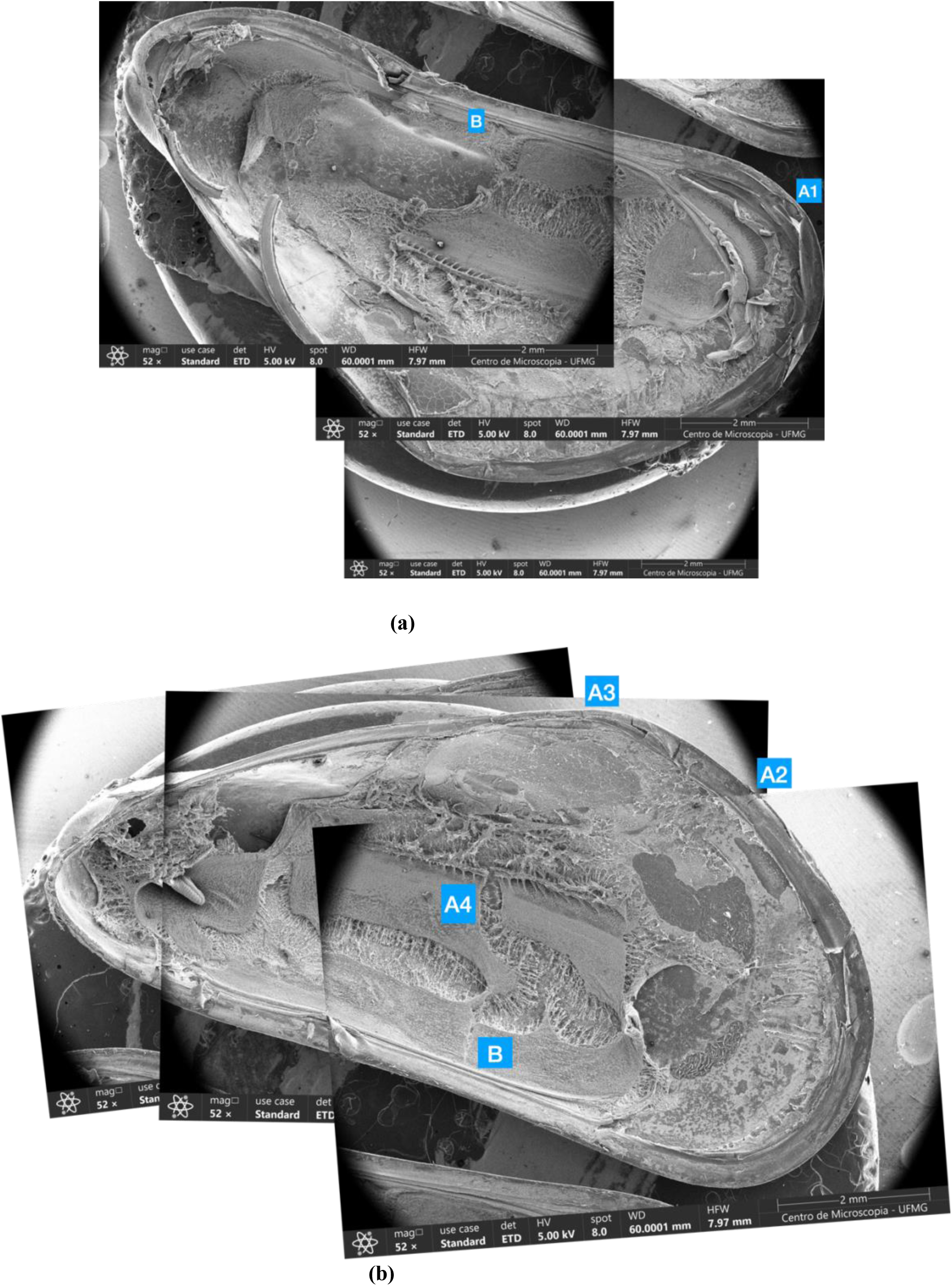
**(a) and 1. (b)** – Two L fortunei shells with parts of the mantle attached. Image montage to allow observation of **regions A1, A2, A3 and A4** as described in the text. The periostracum covers the edge of the shell and **regions A1, A2 and A3** are fractures in this layer that enabled the investigation of the underlying biomineralized layer. The hinge that joins the two shells appears partially detached from the edge of the shell on the underside of the mount in **1. (a)**. Region **A4** is the site of tears in the mantle face. Just above **A4**, structures called mantle edges are observed on both shells. Of the three mantle folds (inner, middle, and outer) we can observe the inner fold, which appears clearly on both shells and is marked with the **letter B**. The periostracum is excreted in the external fold, which appears in the image as a ledge adjacent and parallel to the internal fold. The inner face of the mantle is intact in some areas, torn in other areas where it is possible to see inside and the internal side of the outer face of the mantle in contact with the shell. In some areas it is the biomineralized surface of the shell itself that appears.

Below we describe each of these regions of the shell edge and the results we found.

### 3.2. Shell edge A1

The **A1** region is at the inner end of the L fortunei shell (in detail in the images in Figure 2). The tips are the regions that record the last growth of the bivalve. The images in Figure 2 use two types of SEM sensors, BSE/ABS (backscattered electron/annular backscatter) to observe the surface and its topography (Figure 2.a) and ETD (Everhart-Thornley detector) to obtain images (2.c) generated by secondary electrons coming from deeper regions of the beam/sample interaction.

**Figure 2.**
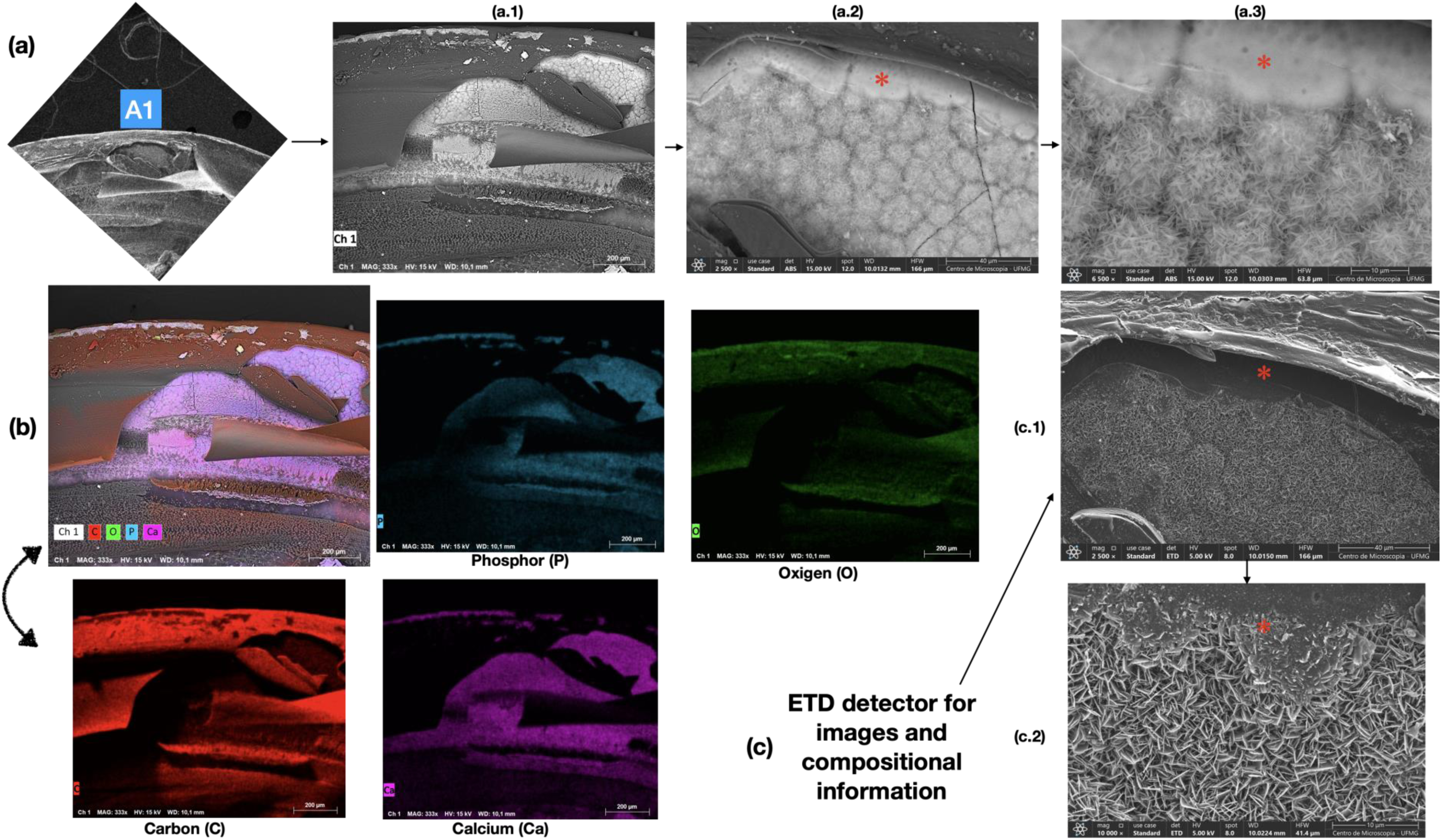
– **(a)** SEM images of region **A1**, in the sequence of images a.1, a.2 and a.3 focusing on the region with aggregates immediately below the periostracum of L. fortunei; **(b)** The image on the right uses energy dispersive spectroscopy (EDS) to highlight the main composition of the region with maps of the main chemical elements present: carbon (C) throughout the periostracum region, calcium (Ca) and phosphorus (P) marking the aggregates and oxygen (O) both in the periostracum and in the aggregates. **(c)** Images c.1 and c.2 were taken with the ETD detector and provide another view of the aggregates of some type of calcium phosphate (P, O, Ca). Image c.2 clearly shows two-dimensional growth in the form of plates. Images a.2 and c.1 were taken at the same magnification as a.3 and c.2. The red asterisks (a.2, a.3, c.1 and c.2) show that the upper surface of these morphologies appears to be coalescing. The same situation can be seen in the next figure 3.d where we have also added a red asterisk to indicate the similarity of the situations.

The periostracum appears on the inner face at the tip of the shell, as it has been secreted by the mantle (seen at the bottom of the image in Fig. 2.a.1). The image of region A1, at the tip of the inner part of the shell, is reproduced at the beginning of the sequence of images in Fig. 2.a, which focuses on the region with aggregates immediately below the periostracum of L. fortunei, at increasing magnifications (from a.1 to a.3). In Fig. 2.a.1 it is possible to see these aggregates at the outermost tip of the shell, indicating biomineralization to the end of the shell. Figures 2.a.2 and 2.a.3 show the morphology of these aggregates more clearly. Figure 2.b shows the maps of the main chemical elements present in this region. Energy dispersive spectroscopy (EDS) obtains semi-quantitative chemical information from the characteristic X-rays of the chemical elements present in the region of incidence of the electron beam. Carbon (C) clearly marks the periostracum, which is an organic film secreted by the folding of the mantle (bottom of fig 2.b, shrinkage due to the drying of the sample caused it to detach from the mantle edge). Oxygen marks both the periostracum and the biomineralized aggregates. The main elements in the aggregates are phosphorus (P), calcium (Ca) and oxygen (O), indicating that they are some kind of calcium phosphate (P,O,Ca). The aggregates appear well individualized in the higher magnification fig 2.a.3. The images using the ETD sensor in Fig. 2.c show the structure of individualized aggregates in detail and, especially Fig. 2.c.2, show growth in the form of plates that grow perpendicular to the image plane. These plates seem to grow by adding nanometer-sized spheroidic particles (Figure 3).

**Figure 3.**
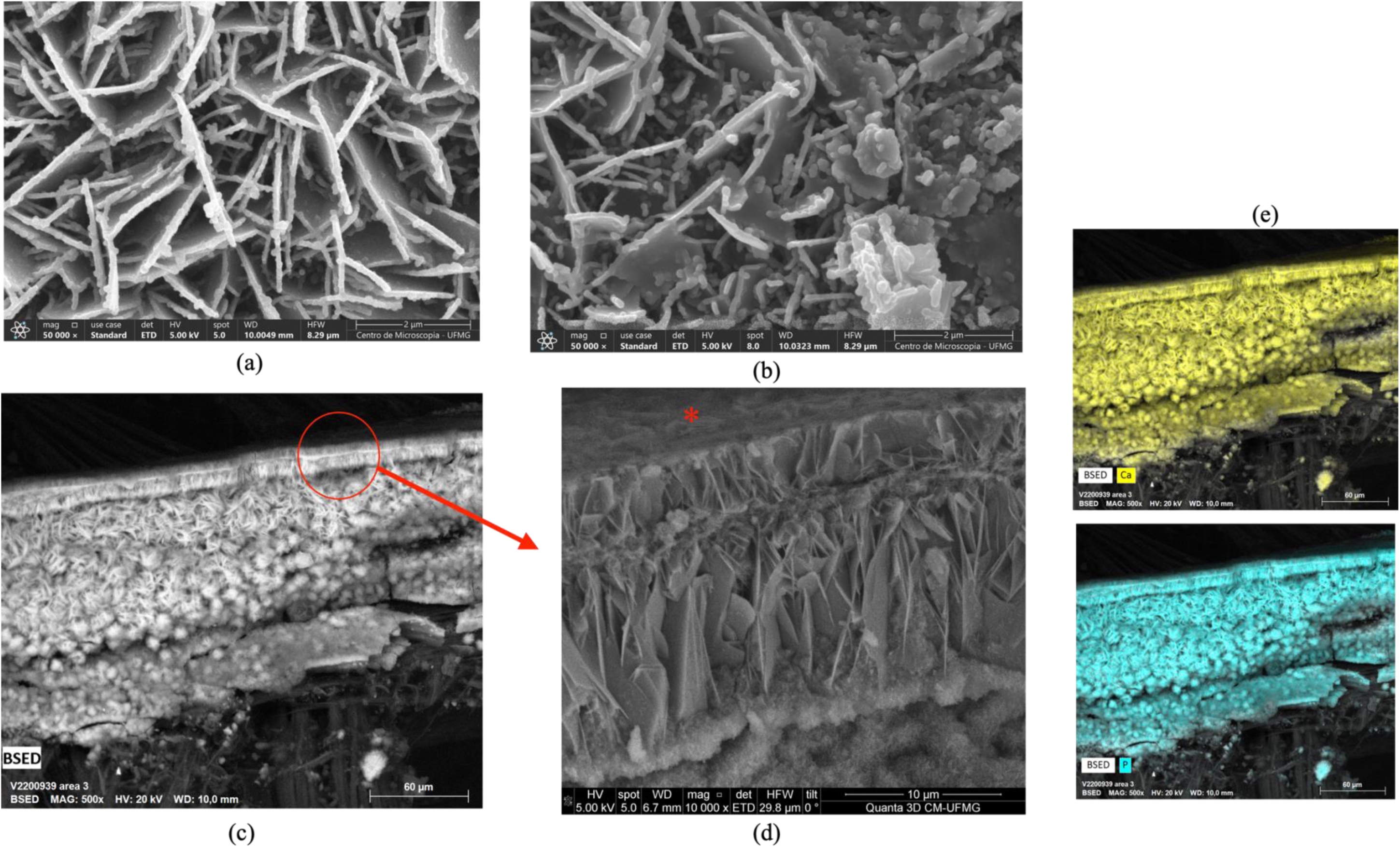
– **(a) and (b)**. Detail of the morphology of the aggregates in the shell tip of L fortunei. Spherical particles of nanometric size are observed in both images. In **(c)** is shown an image taken at the shell tip and the detail in **(d)** shows in lateral view the same plates shown in frontal view in **(a)**. **(e)** shows the calcium (Ca) and phosphorus (P) maps marking the same region as the aggregates shown in **(c)**. The **red asterisk** in **(d)** is to show that the surface above the plates appears coalesced. See images in 2.a.2, a.3, c.1 and c.2 where this coalesced surface also appears above the plates. Images in **(a)** and **(b)** are from one L fortunei specimen while figs. **(c)**, **(d)** and **(e)** are from another L fortunei specimen.

In both Fig.3.and Fig.3.b it is possible to see a two-dimensional growth of plates of a phosphate compound. These plates grow due to the adhesion of phosphate nanoparticles. The interaction between the plates seems to indicate that the surface of one plate causes nucleation and therefore the appearance of another plate. The nanometric spheres are possibly amorphous and may crystallize when they organize themselves into plates.

### 3.3. Shell edge A2

**Area A2** is in the shell shown in Fig. 1.b and is shown in detail in the images in Figure 4. The sequence of images a.1, a.2 and a.3 focus on the region of aggregates immediately below the periostracum of L. fortunei. Aggregates whose composition estimated by EDS spectroscopy (Fig. 4.b.1) indicated the presence of phosphorus (P), oxygen (O) and calcium (Ca) are observed immediately below the periostracum. The spectrum indicating the concentration of these elements is shown in Fig. 4.b.2. The maps of the distribution of these chemical elements are shown in the right part of the figure. Carbon is marked throughout the periostracum, calcium and phosphorus mark the aggregates, and oxygen is marked both in the periostracum and in the aggregates. The (semi-quantitative) values for the concentrations of the elements are given in Table 1.

**Figure 4.**
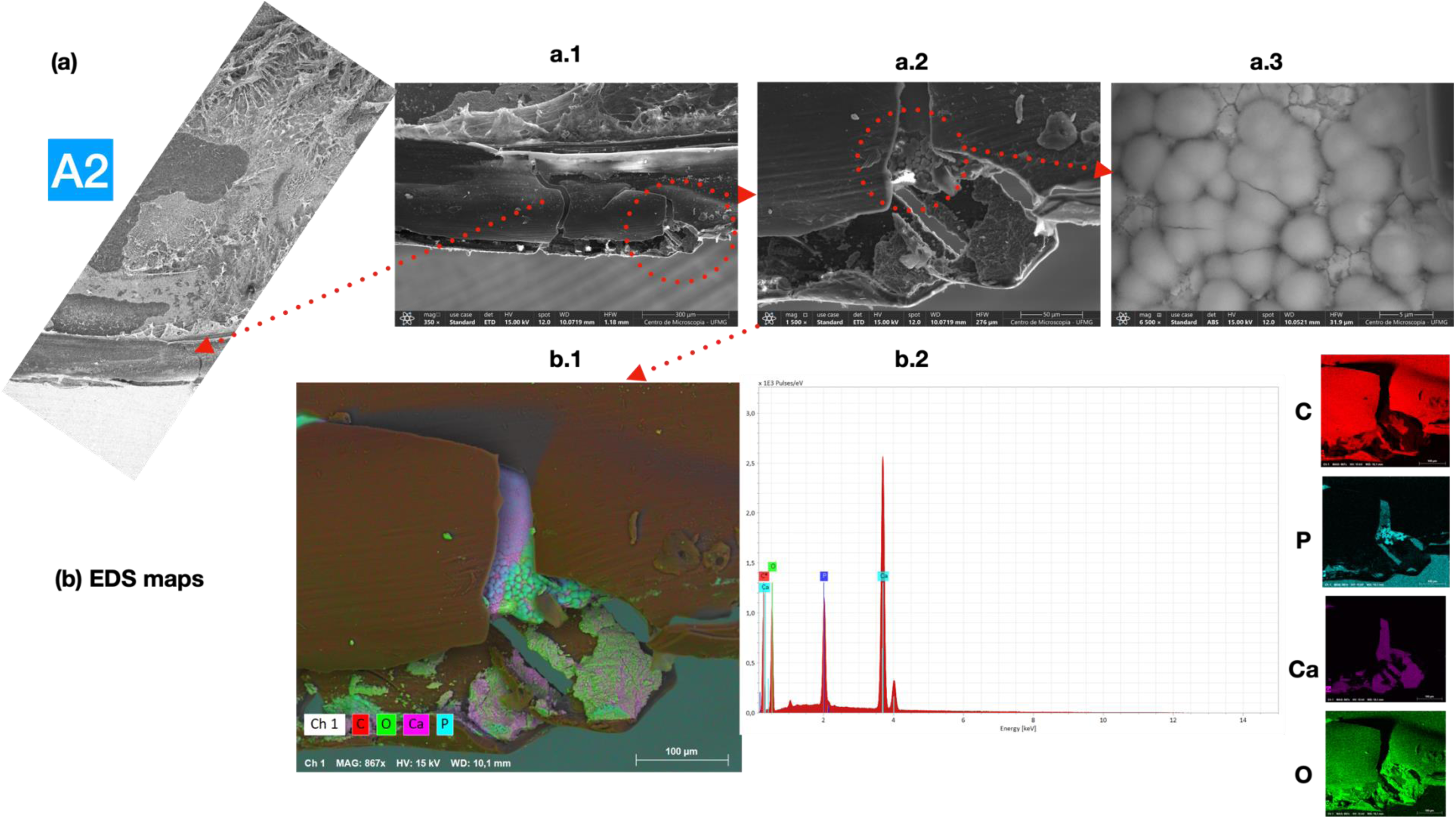
– **(a)** SEM images of region **A2**, in the sequence of images a1, a2, and a3, focusing on the aggregates immediately below the periostracum of L. fortunei. **(b)** Image b.1 shows the main composition of the region with maps of the main chemical elements present in the right part of the figure: carbon (C) throughout the periostracum region, calcium (Ca) and phosphorus (P) marking the aggregates, and oxygen (O) in both the periostracum and the aggregates. **Table 1** shows the concentration of the main elements present, as depicted in b.2.

**Table 1.**
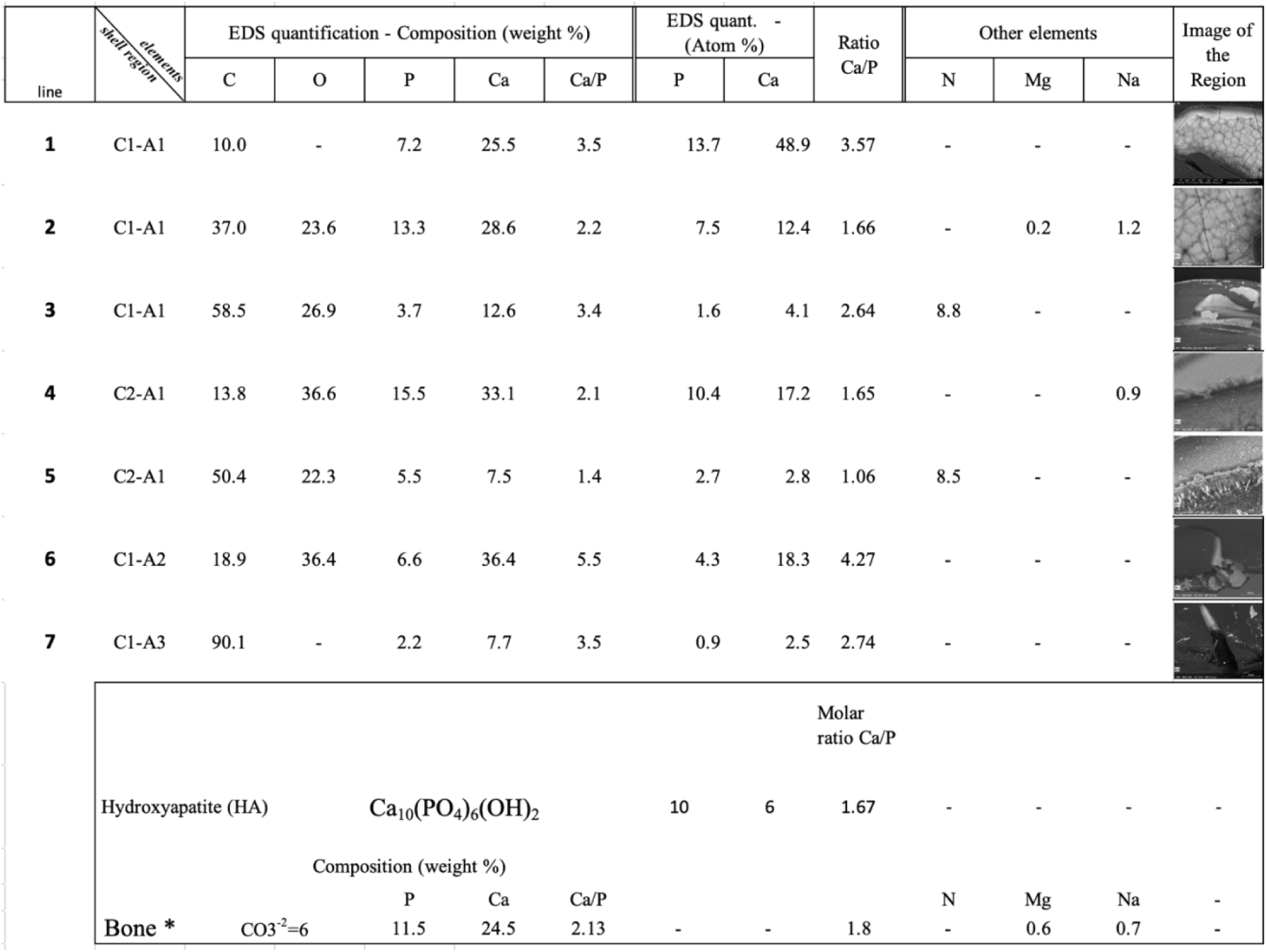
Elemental composition (% by weight, semi-quantitative) of regions A1, A2 and A3 of the edge of the L fortunei shell (images in the last column) obtained by EDS spectroscopy, Ca/P ratio in % by weight and % molar. The lower part of the table shows the chemical composition, Ca/P ratios in weight and molar, of the carbonated hydroxyapatite from bone proposed by LeGeros (44) (marked with a black asterisk) and the molar ratio of hydroxyapatite.

The images in Figure 5 show the morphology of the aggregates depicted in the previous figure. Immediately below the periostracum are aggregates marked by EDS (Fig. 4.b.1) as consisting of phosphorus (P), oxygen (O) and calcium (Ca). These aggregates (seen in detail in Fig. 5.b.1) tend to form a complete layer by coalescence (Fig. 5.a.1 and 5.a.2). The structural organization of these aggregates, whether crystalline or amorphous, could not be characterized. Immediately below these aggregates appears what looks like the nacreous layer of the L. fortunei shell, shown in Fig. 5.a.3. This nacreous layer is strongly marked as calcium (Ca) in Fig. 4.b.1 but seems not to contain phosphorous (P). Finally, nanometric particles are seen in Figures 5.a.2 and 5.b.3 and are circled in red. The origin (whether detached from or attached to the aggregates), direction (downward, sideways, etc.) and chemical composition of these nanoparticles is unknown.

**Figure 5.**
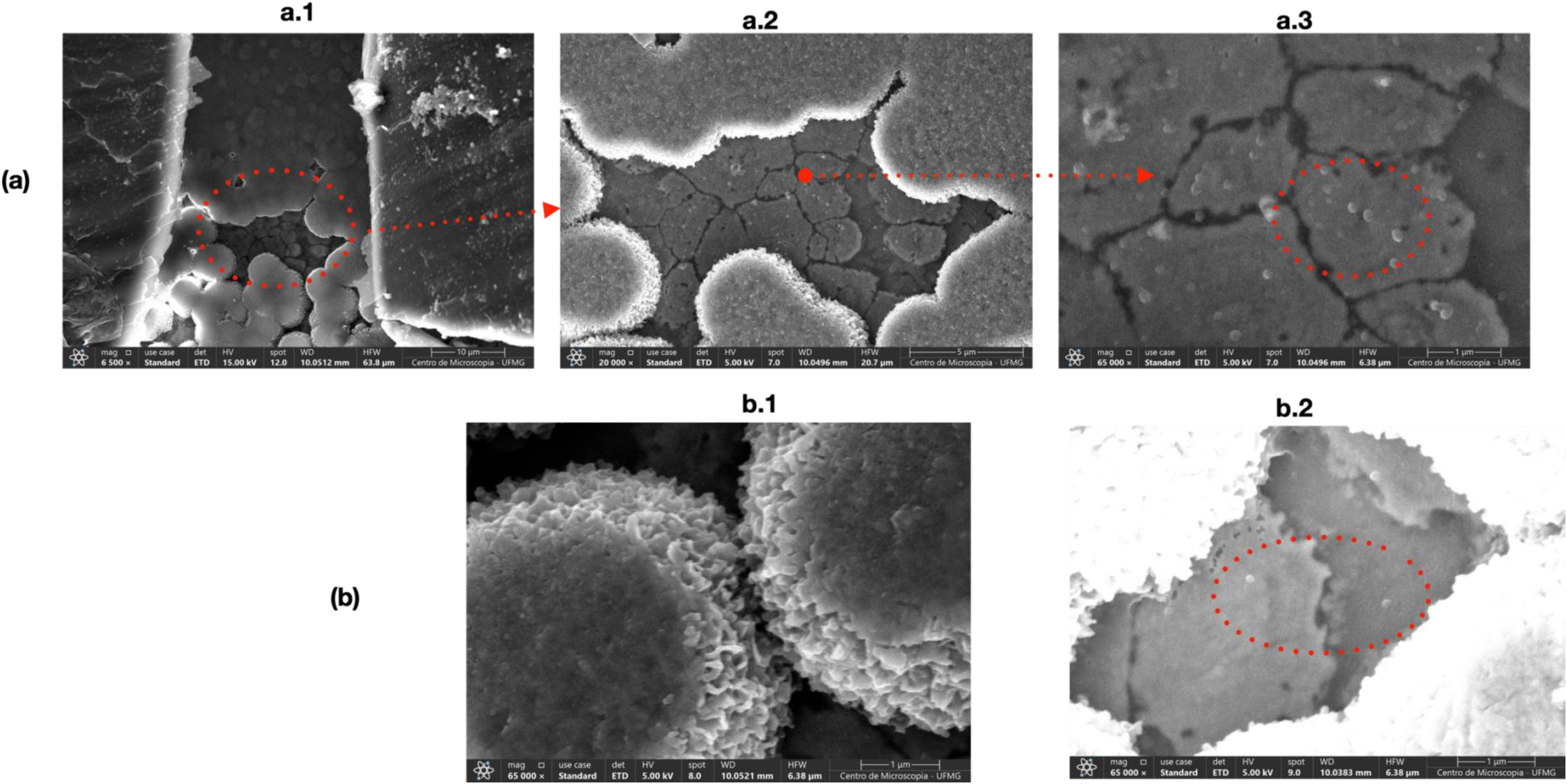
– **(a) and (b)**. Detailed view of the morphology of the aggregates in **region A2**, as depicted in the previous figure. Immediately below the periostracum, aggregates marked by EDS with compounds of phosphorus (P), oxygen (O), and calcium (Ca) are present (Fig. 4.b.1). These aggregates (seen in detail in b.1) tend to form a complete layer through coalescence (**Figs. 5.a.1 and 5.a.2**). Immediately below these aggregates, the nacreous layer appears, as shown in image **a.3**. Nanoparticles are visible in Figures 5.a.3 and 5.b.2, circled in red. The origin, direction (downwards, sideways, etc.), and chemical composition of these nanoparticles are unknown.

### 3.4. Shell edge A3

**Region A3** has two fractures on the edge of the shell (circled in red in the A3 image shown in Figure 6). Figure 6.a.1 shows energy-dispersive spectroscopy (EDS) results to highlight the main composition of the region, with the elemental maps (6.a.2) showing carbon throughout the periostracum region and calcium and phosphorus marking the aggregates. The concentrations (semi-quantitative) of these elements are presented in Table 1.

**Figure 6.**
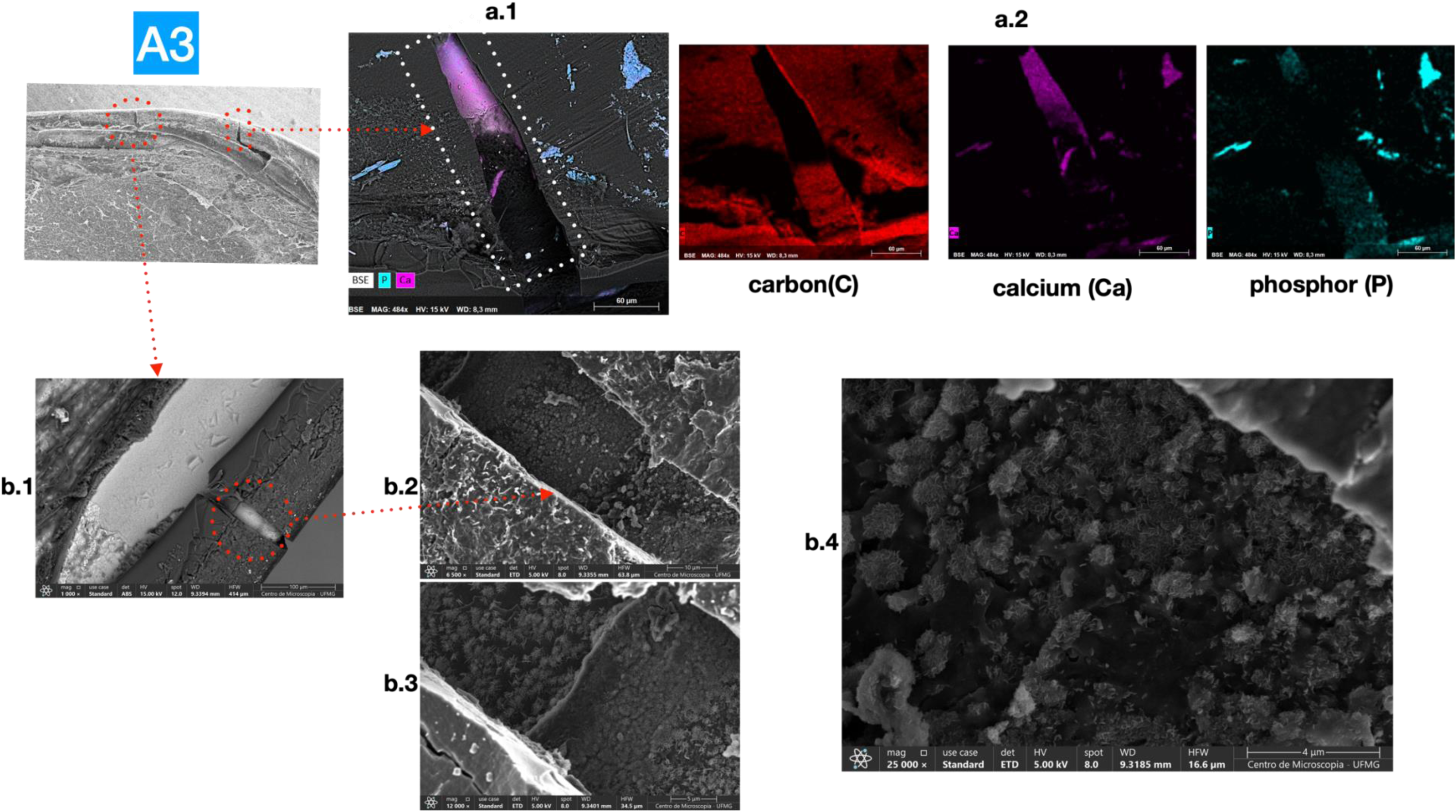
– **(a)** shows SEM images of **region A3**, where two fractures are present on the edge of the shell (circled in red). Image in Fig.6. a.1 is an energy dispersive spectroscopy (EDS) image that highlights the main composition of the region, with maps (Fig. 6.a.2) of the chemical elements carbon, calcium, and phosphorus marking the aggregates throughout the periostracum region. The semi-quantitative concentrations of these elements are presented in Table 1. **(b)** The sequence of images in Figs. 6.b.1, b.2, b.3, and b.4 focus on the fracture of the leftmost shell edge. The aggregates containing P, O, and Ca have the same morphology as the **A.1 region** (see above Fig. 2 and Fig.3). In image of Fig. 6.b.4, it is possible to observe that these aggregates tend to coalesce (see above red asterisk in Fig. 2.a.2 and Fig. 3.d).

Images 6.b.1 through 6.b.4 focus on the fracture of the leftmost edge. The aggregates containing P, O, and Ca have the same morphology as those in region A.1 shown in Fig. 3. Image in Fig. 6.b.4 shows that these aggregates tend to coalesce and form continuous plates, as also observed in the image presented in Fig. 3.b.

### 3.5. Mantle-nacre and region **A4**

In addition to the edges, we investigated areas where layers of the mantle were directly above the nacre, as shown in the SEM images in Figure 7 (a.1, a.2, and a.3). Image 7.a.4 was taken with the T1 detector (equipment Apreo-2C) to highlight the topography and the difference between the mantle and the nacreous layer of the shell. The chemical element maps (below 7.a.4) show a continuous band composed of phosphorus (P) and calcium (Ca). The thickness of the phosphate layer, measured in the SEM, is approximately 3 μm (see Figure S6 Supplementary Material). The calcium phosphate layer is composed of nanoparticles of varying sizes, as shown in the higher magnification image in 7.a.3. Below this layer lies the nacreous layer, which is composed of calcium carbonate. In Fig. 5, calcium phosphate aggregates are also associated with nanoparticles, and just below them, the nacreous layer is already visible.

**Figure 7.**
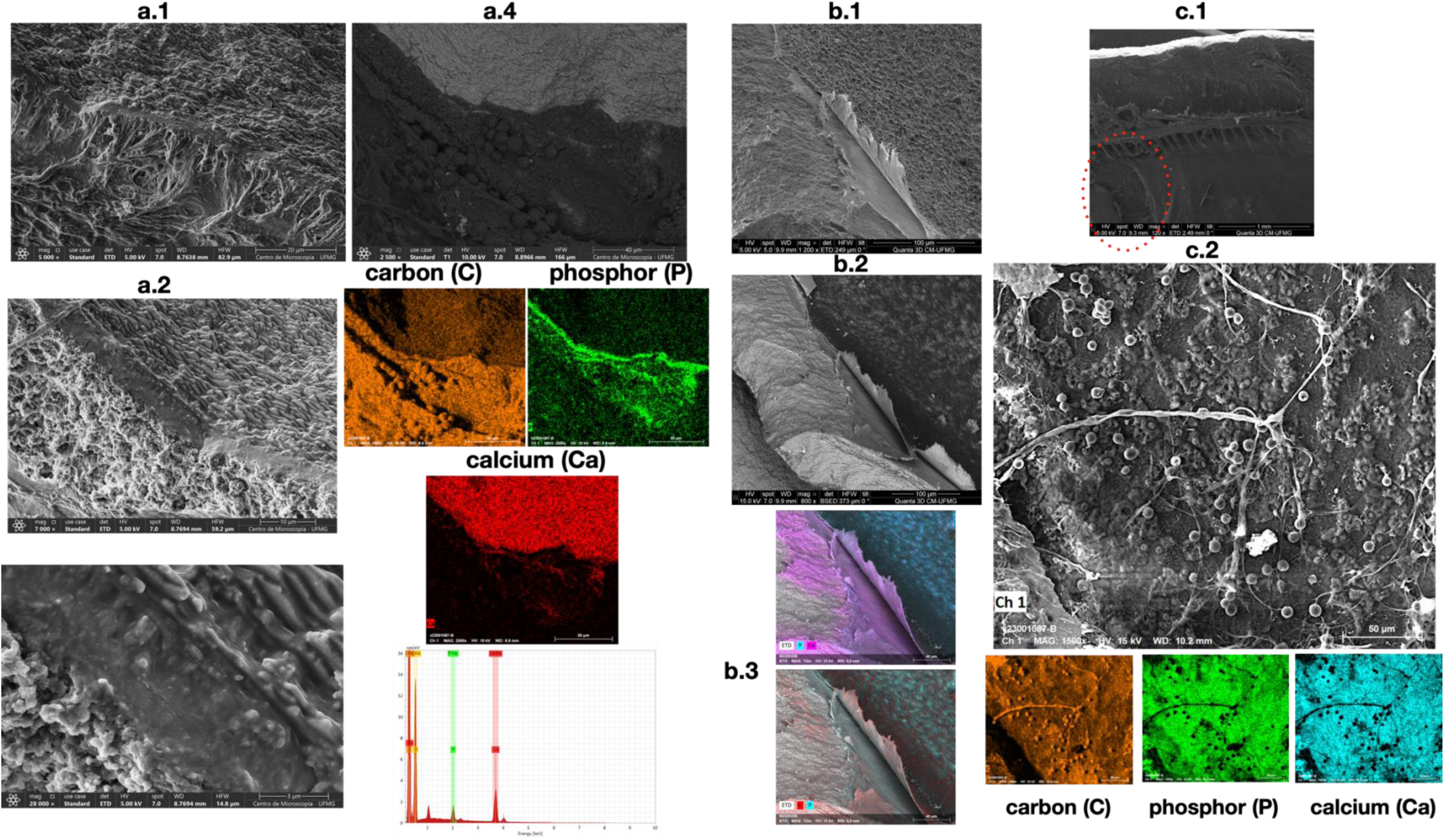
– **(a)** SEM images of regions where the mantle layer overlaps with the nacreous layer of the L. fortunei shell (a.1, a.2, and a.3). Image a.4 highlights the topography and the difference between the mantle and the nacreous layer of the shell, taken with the T1 detector. The chemical element maps (below a.4) show a continuous band of phosphorus (P) and calcium (Ca). Carbon (C) clearly outlines the mantle with spheres of mucus. **(b)** Images b.1 to b.3 illustrate the presence of P, Ca, and C in the nacre-mantle interface. **(c) Region A4** (see also Fig. 1.b above) is a place where the inner face of the mantle is torn, and it is possible to investigate the *internal side* of the mantle outer layer in contact with the L. fortunei shell. The map of carbon outlines various objects such as inner mantle fibers and possibly mucus balls. Additionally, phosphorus and calcium are present throughout the inner surface of the mantle. Image c.1 shows an area of the shell containing the mantle with part of the bivalve’s musculature. Torn regions of the mantle (red outline in c.1) are shown in detail in c.2. This image c.2 shows the mantle layer in contact with the shell.

Figure 7.a.4 displays the carbon map (C) which outlines the mantle layer. The presence of spheres, potentially mucus, can be observed. Additionally, fibers that make up the mantle’s internal structure are visible. Images 7.b.1 to b.3, taken from another shell sample, also show the presence of P, Ca, and C at the nacre-mantle interface.

Image 7.c.1 depicts the inner surface of the mantle and a structure known as the mantle rim (42,43) of the bivalve. Due to tissue dehydration, the mantle edge appears near the center of the shell. Detailed images of torn mantle areas (outlined in red in 7.c.1) reveal the inner layer of the mantle in contact with the shell. Region A4 (Fig. 1.b) is one such area where the torn mantle allows for investigation of the side in contact with the L fortunei shell. The diagram in image 7.c.2 shows the distribution of carbon (C) in the internal mantle fibers, mucus sphere, and in other forms. Additionally, phosphorus (P) and calcium (Ca) are present throughout the inner surface of the mantle that is in contact with the shell.

### 3.6. **Region B**: Mantle Edge and Mantle Surface

The edges of the mantle fold are essential regions for understanding the process of biomineralization. The process begins with the release of a film called periostracum, which is composed of chitin, collagen, and proteins. This film is released by the outer fold of the mantle and is formed by adhered films (2 layers). Understanding this process is crucial to understanding biomineralization. The excretion by the cells located in the outer fold, its polymerization resulting in a hydrophobic film, and its adhesion to the existing periostracum are still not well understood. Two additional folds, namely the intermediate and internal folds, have been identified. These structures are visible in the SEM images of the mantle folds (Figures 8.a.1, 8.a.2, and 8.a.4). Image of Fig. 8.a.1 reveals that the folds are regions with a high density of cilia, which is better visualized in 8.a.2. Additionally, 8.a.4 shows the large quantity of mucus spheres produced in this fold. The EDS spectroscopy measurements in 8.a.3 display-the chemical maps, indicating that carbon (C) and oxygen (O) outline the entire area of the cilia. Carbon marks and delimits the image of the mucus spheres. However, the presence of phosphorus and calcium in the EDS spectrum shown in 8.a.3 does not provide specific identification for any object due to their low concentrations.

**Figure 8.**
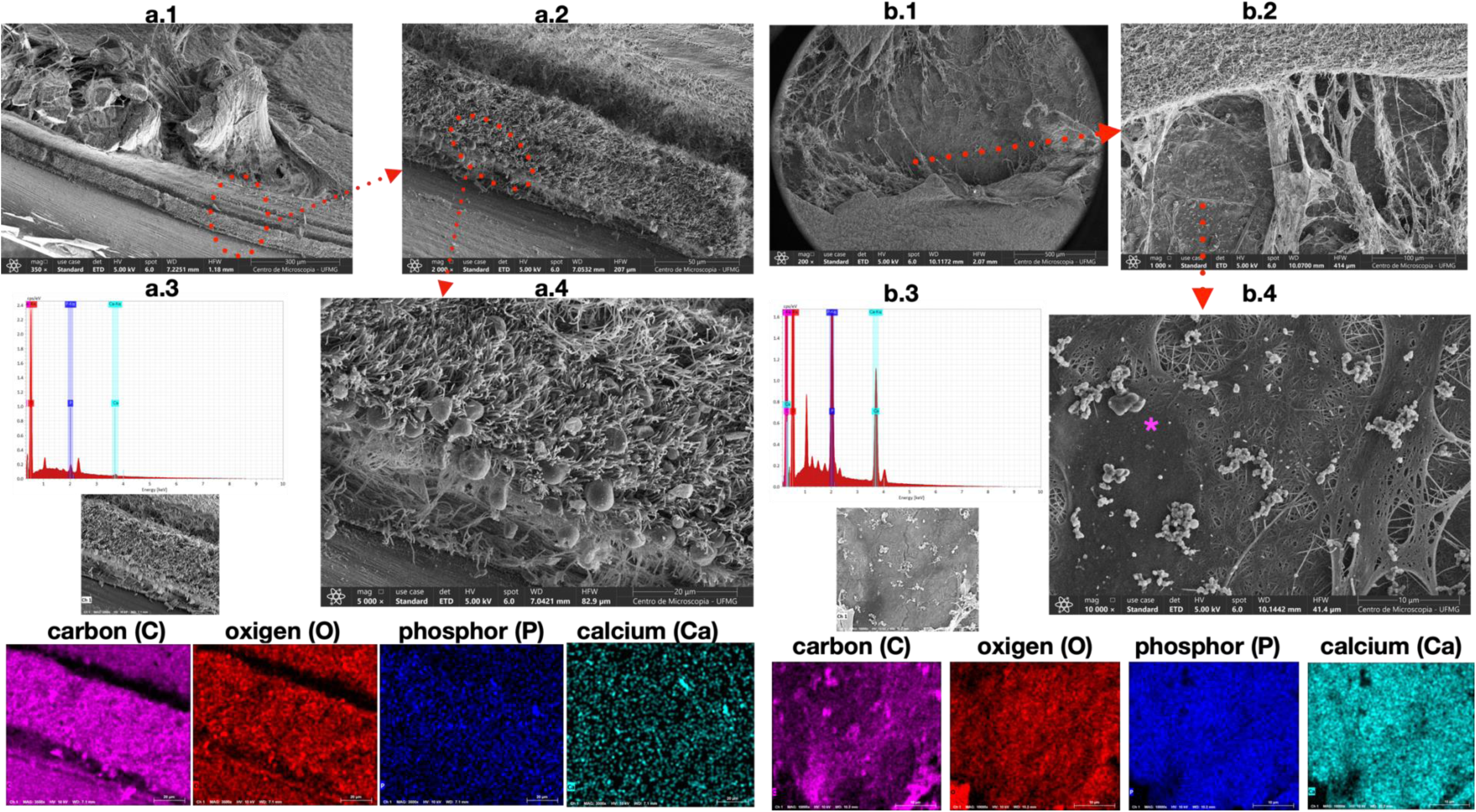
– **(a)** Scanning electron microscope (SEM) images of the mantle folds (a.1, a.2, and a.4). The folds are regions with a high density of cilia, as seen in a.2. In a.4, a large quantity of mucus spheres produced in this fold can be seen. The EDS spectroscopy in a.3 and the chemical maps presented indicate that carbon (C) and oxygen (O) outline the area, while the presence of phosphorus (P) and calcium (Ca) appears to be very low. **(b)** SEM images of the mantle’s surface and interior are shown in b.1, b.2, and b.4. The ciliated surface of the mantle is clear visible in b.1 and better visualized in b.2. The large number of cilia present on the surface are believed to aid in capturing micro and nanometric particles in the water. The image in b.4 shows the inner surface of the outer face of the mantle, which is in contact with the shell. Several layers of fibers and aggregates of particles (magenta asterisk) are visible on this surface. The EDS spectroscopy in b.3 and the chemical maps presented indicate that the particle aggregates on the inner surface of the outer face of the mantle are marked with carbon. The concentration of phosphorus, oxygen, and calcium is significant compared to the spectrum in a.3.

SEM images 8.b.1, 8.b.2, and 8.b.4 show the internal surface of the mantle, which houses the viscera of L. fortunei. The ciliated surface is visible in 8.b.1 and better visualized in 8.b.2. The numerous cilia are likely crucial for capturing micro and nanometric particles when the shell is open. Figure S.4 in the supplementary material shows that the mantle of L. fortunei has ciliated areas surrounding non-ciliated areas where particles may interact with the mantle wall.

Additionally, image 8.b.4 displays the inner surface of the outer face of the mantle, which is in contact with the shell. This surface exhibits several layers of fibers and aggregates of particles. The EDS spectroscopy in section 8.b.3 and the chemical maps indicate that the particle aggregates on the inner surface of the outer face of the mantle are composed of carbon (C). The concentrations of phosphorus (P), oxygen (O), and calcium (Ca) are significantly higher compared to the spectrum in section 8.a.3.

### 3.7. Quantification of the Chemical Composition of Shell Edge Regions by EDS and Ca/P Ratio

In the shell edge fractures shown in the previous figures, where the presence of distinct morphologies for phosphate compounds was identified, quantifications of the elements carbon (C), oxygen (O), phosphorus (P) and calcium (Ca) were carried out by EDS spectroscopy. Table 1 shows these quantifications in the typical microstructures of regions of the edge of the L fortunei shell, designated A1, A2 and A3 and already shown in Figure 1 and subsequent figures. The images are of two L fortunei individuals, designated C1 and C2 for simplicity. Rows 1, 2, and 3 of the table are from edge A1 of the shell sample from individual L fortunei C1. Rows 4 and 5 are from the shell sample of individual C2. Row 6 shows results from area A2 of shell sample C1. Row 7 of the table is from area A3 of the rim of shell sample C1. The images shown in the last column of the table are from the areas where quantification was performed.

The Ca/P ratios (% atomic mass) were calculated from the atomic percentage data of the elements quantified by EDS (rows 1 to 7 of Table 1). At the bottom of the table are the molar ratios for hydroxyapatite (HA), 1.67, and carbonated hydroxyapatite (1.8). Some Ca/P (atomic mass) ratios obtained from EDS spectroscopy (rows 2 and 4) show values close to the HA molar ratio (1.67), but the other values for the Ca/P ratio are very different from HA and carbonated HA. The highest value for the Ca/P ratio occurs in the A2 region hypothetically due to the more advanced stages of the phosphate to carbonate substitution. Some quantifications of magnesium (Mg) and sodium (Na) have also been included to show that these elements are always present in much lower concentrations. The quantification of nitrogen (N) has been included in the table for some morphologies to reflect its origin, as well as zinc (Zn), shown in figure S7 in the Supplementary Material.

### 3.8. Quantification of the chemical composition of L. fortunei and P. perna shells by ICP-OES Spectroscopy

Table 2 shows the concentrations of various metals and non-metals in the shell of the freshwater golden mussel (Limnoperna fortunei). The concentration of calcium is very high as it is the main component of the shell. Ba, Fe, Mg, Sr, Na and Zn reach concentrations of fractions of a percent. These metals and other traces of heavy metals, present in both fresh and marine water, were assimilated in the mantle, in the gills or absorbed by the digestive system. Sodium (Na) reaches decimal percentages, probably because it is essential for most metabolic processes. The concentrations of other metallic elements are very close to the values obtained in other studies (45,46), especially for calcium ions, since CaCO_3_ represents more than 95% of the composition of the shell.

**Table 2.** Metal and non-metal element composition of the shells of Limnoperna fortunei and Perna perna derived using ICP-OES spectroscopy (in %)

| Sample | Year | Ba | Ca | Fe | P | Mg<br>% | Na | Sr | Zn | K |
| --- | --- | --- | --- | --- | --- | --- | --- | --- | --- | --- |
| *Limnoperna fortunei* Shell | 2016 | 0.0159 | 35.08 | 0.040 | - | 0.0235 | 0.220 | 0.059 | 0.0090 | 0.0032 |
| *Limnoperna fortunei* Shell | 2022 | 0.0161 | 36.07 | 0.011 | - | 0.0180 | 0.537 | 0.120 | 0.0003 | 0.0037 |
| *Limnoperna fortunei* Shell | 2023 | 0.0002 | 38.71 | 0.002 | 0.026 | 0.0190 | 0.640 | 0.120 | 0.0003 | 0.0190 |
| *Perna perna* Shell | 2022 | 0.0003 | 38.64 | 0.002 | - | 0.0159 | 0.663 | 0.105 | <2 | 0.0133 |
| *Perna perna* Shell | 2023 | 0.0340 | 39.05 | 0.043 | 0.039 | 0.0230 | 0.250 | 0.071 | 0.0008 | 0.0029 |

The concentration of phosphorus (P) is in the order of ppm in the whole shell and differs greatly from the values found at the edges of the shell growth shown in Table 1. Studies that include the chemical composition of the shell do not report the presence of phosphorus, possibly because it happens at very low concentration (47).

### 3.9. Quantification of the chemical composition of L. fortunei and P. perna shells by WDXRF spectroscopy

Table 3 shows the concentrations of different metals and non-metals present in the shells of L. fortunei (freshwater) and P. perna (marine) obtained by wavelength dispersive fluorescence spectroscopy (WDXRF). These are semi-quantitative values, but they are in excellent agreement with the concentrations obtained using ICP-OES. As the sample was a whole shell, the phosphorus concentrations are in the order of ppm (parts per million). Therefore, the occurrence of calcium phosphate should be limited to the edges of growth (in area and in thickness) and only occur there. The calcium concentration is >98% for both the shell of the freshwater L fortunei and the marine P perna bivalves. The concentrations of the other chemical elements are less than 2%.

**Table 3.** Metal and non-metal element composition of the shells of Limnoperna fortunei and Perna perna derived using WDXRF spectroscopy (in %)

| Sample | Year | Al<br>% | Ba | Ca | Cl | Fe | K | Mg | Na | P | S | Si | Sr |
| --- | --- | --- | --- | --- | --- | --- | --- | --- | --- | --- | --- | --- | --- |
| *Limnoperna fortunei*<br>Shell | 2023 | 0.051 | - | 98.190 | 0.280 | 0.033 | 0.044 | 0.050 | 0.760 | 0.037 | 0.240 | 0.120 | 0.195 |
| *Perna perna*<br>Shell | 2023 | 0.110 | 0.490 | 98.100 | 0.420 | 0.230 | - | 0.062 | 0.430 | 0.089 | 0.210 | 0.140 | 0.120 |

### 3.9. X-ray diffraction (XRD) of the shell tips of L fortunei and P perna

Figure 9 shows the diffractograms of the shells of L fortunei and P perna. As can be seen, the diffractograms are practically identical, indicating the presence of the aragonite and calcite phases. Aragonite is the most abundant phase in the two shells. The diffractogram indicates a ratio of (aragonite)10:1 (calcite), considering the height of the peaks located at the 2θ angle of 46° for aragonite and 40° for calcite (48).

**Figure 9.**
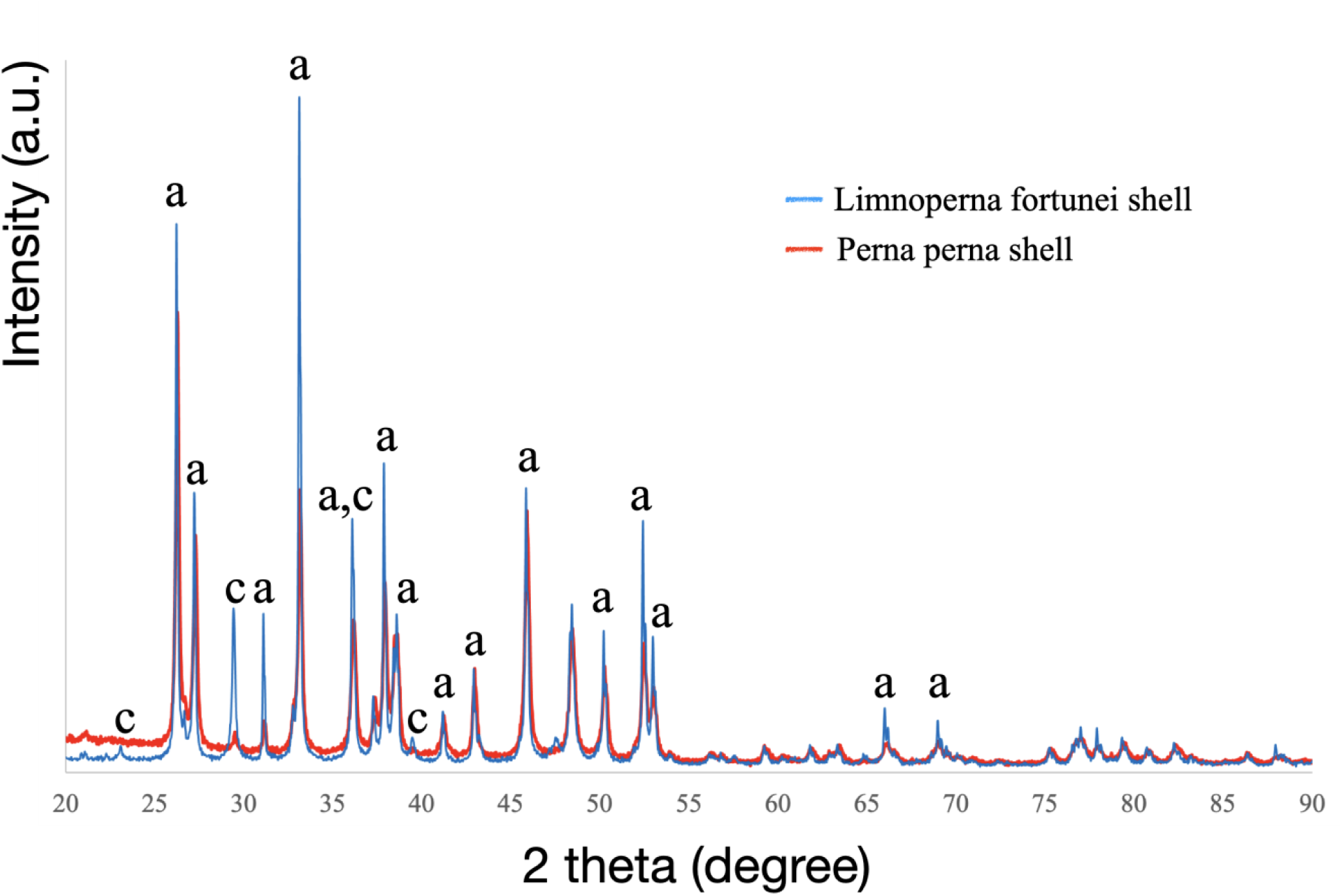
– X-ray diffractograms of mussel shells. The aragonite and calcite phases are observed in the XRD diffractograms of L. fortunei and P. perna. Calcium phosphate phases were not detected. a: aragonite, c: calcite.

No other phase was detected, in agreement with other results (ICP-OES and WDXRF) that prove a very low concentration of phosphorus (P) in the shell. This makes apatite, hydroxyapatite or other P phase detection difficult by XRD. However, we were not able to identify the 2θ peak at 26.21°, which could be one of the main peaks of hydroxyapatite [25.9° (002)](49). It should also be noted that the crystalline phases of calcium phosphate, especially hydroxyapatite, have their main diffraction peaks coinciding with aragonite and calcite, which hardens detection.

### 3.10. FTIR and FTIR-ATR spectroscopy of the shell tips of L fortunei and P perna

Figure 10 shows the FTIR and FTIR-ATR results (upper part of the figure) for shell tip samples of L. fortunei and P. perna where the main functional groups could be identified. Carbonate is observed in all spectra (FTIR and FTIR-ATR) of L. fortunei and P. perna. It can be attributed to the absorption bands at 700, 712, 862, 1084, and 1466 cm^-1^ (50). The band at 2924 cm^-1^ is probably due to the presence of chitosan in the shell and in the periostracum attached to it (51). The bands between 3000-3600 cm^-1^ are attributed to hydroxyl groups and adsorbed water. In FTIR-ATR these features were not observed. The absorption bands at 2359 and 1643 cm^-1^ could be attributed to the antisymmetric stretching of CO_2_ absorbed from air(52,53).

**Figure 10.**
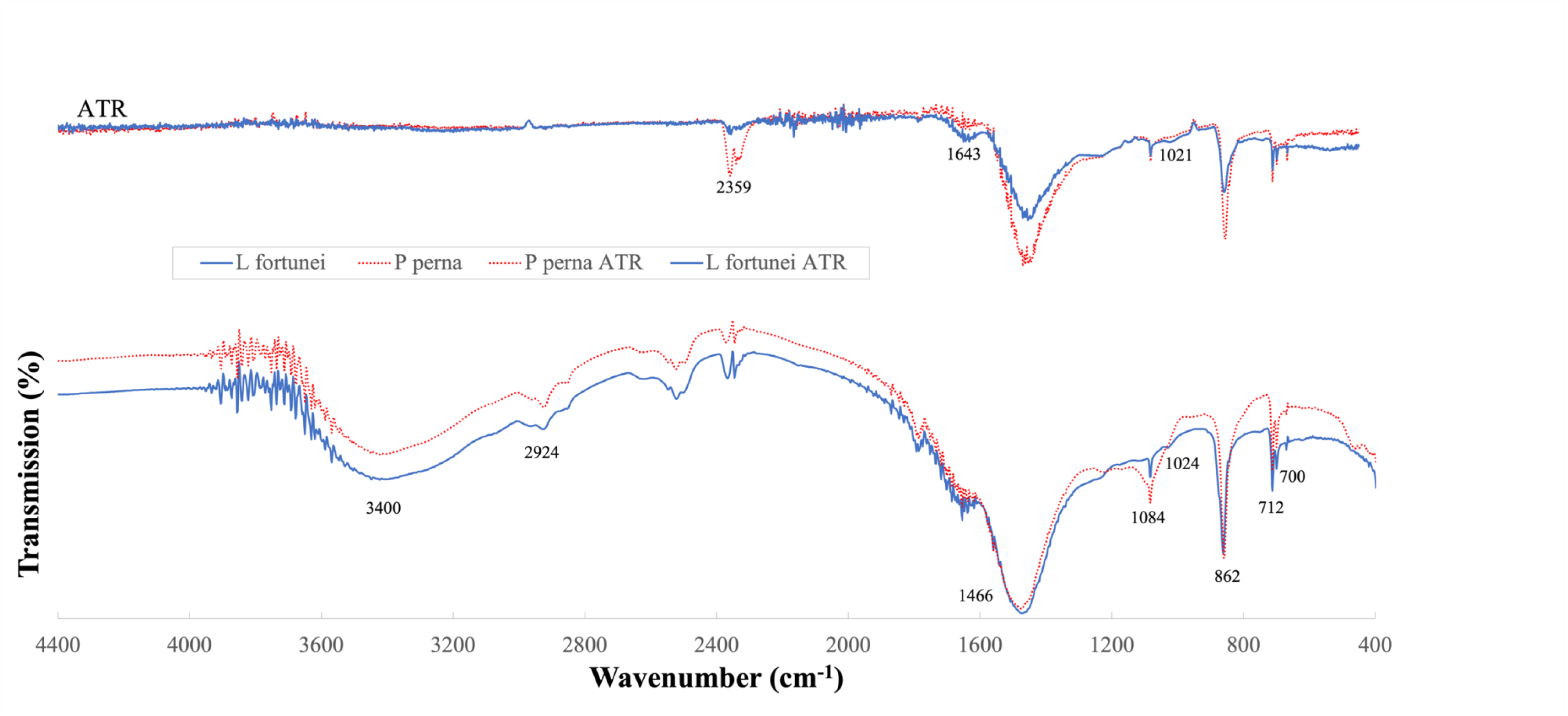
– FTIR and ATR-FTIR spectra (upper part of the figure) of mussel shells. The aragonite phase is observed in all spectra (FTIR and FTIR-ATR) of L. fortunei and P. perna by the absorption bands at 700, 712, 862, 1084 and 1466 cm^-1^. The band at 2924 cm^-1^ can be attributed to the chitosan (51) present in the shell. The band from 3000 to 3600 cm^-1^ is associated with the vibration of hydroxyl (OH^-^), but also with adsorbed water. This band does not appear in the FTIR-ATR spectrum. The bands at 1643 and 2359 cm^-1^ as well as other bands not marked in the FTIR spectrum can be attributed to carbonate(52,53). The band at 1024 in the FTIR of the shells of L fortunei and P perna can be attributed to the chemical group phosphate (PO_4_^-3^) (54,55)

The chemical group phosphate appears as the very weak and broad absorption bands at 1024 (FTIR) and 1021 cm^-1^ (FTIR-ATR) in the shells of L fortunei and P perna (54,55). Other absorption bands attributed to the phosphate group may partially or totally superimpose with those associated with the carbonate group and could not be surely identified.

### 3.11. Carbonic Anhydrase Assay

Table 4 presents the results obtained for the carbonic anhydrase assay in demineralized suspensions of L. fortunei and the reference material bovine carbonic anhydrase (BCA).

**Table 4.** – Carbonic Anhydrase Assay.

| column<br>row | 1 | 2 | 3 | 4 | 5 | 6 |
| --- | --- | --- | --- | --- | --- | --- |
|  | sample | T(s).<br>Method 1 | (To-T)/To<br>(WAU unit) -<br>method 1 | T (s)<br>Method 2 | (To-T)/To<br>(WAU unit) -<br>method 2 | Specific activity (WAU units/mg) |
| 1 | Blank (To) | 90 | - | 120 | - | - |
| 2 | BCA | 34 | 0.62 | 70 | 0.42 | Method 1: 2,99 x10<sup>3</sup>,<br>Method 2: 2,4 x 10<sup>3</sup> |
| 3 | DS-AA-Lf | 61 | 0.32 | 70 | 0.42 | - |
| 4 | DS-EDTA-Lf | 120 | - | 120 | - | - |
| 5 | DS-EDTA-Lf<br>(ZnCl<sub>2</sub> added) | 71 | 0.21 | - | - | - |

Method 1: reference (36); Method 2: reference (37). T: time in seconds for the pH drop from 8.3 to 7.3 in method 1 and from pH 8.2 to 6.5 in method 2. To: time for pH drop with buffer solution only. Blank: buffer only. BCA: bovine carbonic anhydrase. DS-AA-Lf: demineralized shell solution in acetic acid. DS-EDTA-Lf: demineralized solution with EDTA. DS-EDTA-Lf (ZnCl<sub>2</sub> added): ZnCl<sub>2</sub> added to the demineralized solution with EDTA. WAU: Wilbur-Anderson Units. The results for blank, BCA, and DS-AA-Lf in method 1 are the average of six independent experiments.

The results indicate that the suspensions obtained by demineralization with acetic acid and EDTA resemble the behavior of BCA when in the presence of excess CO₂, causing a pH drop from >8 to values below 6.5. This result is due to the reaction catalyzed by carbonic anhydrase in the interconversion of CO₂ and water into a bicarbonate ion and a proton:

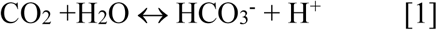

Assays conducted with buffer solutions only (blank) showed much longer times, indicating the absence of the enzymatic catalyst. Assays with demineralized shell suspension with 0.5 M EDTA at pH 8.0, without the subsequent addition of ZnCl₂ (line 4 of Table 4), showed a time interval equal to that of the solution containing only the buffer.

Both methods involve the addition of CO₂ to the suspension. The pH in the buffer suspension also tends to acidify over time due to the increasing CO₂ concentration. However, the times are longer in the absence of carbonic anhydrase, as shown in Table 4, indicating that pH drop is slower without the presence of this enzyme.

The Wilbur-Anderson unit (WAU) value of the reference material (column 3 of Table 4), bovine carbonic anhydrase (BCA), is more than double the values obtained for the demineralized shell suspensions, indicating a higher specific enzymatic activity (column 6 of Table 4). The specific enzymatic activities of the suspensions were not calculated, as the objective of this study was to demonstrate the presence of enzymes in the shell with activity like that of carbonic anhydrase.

## 4. Discussion

The SEM images using backscattered electrons and energy dispersive X-rays (EDS) detectors for mapping of the chemical elements calcium (Ca), carbon (C), phosphorus (P) and oxygen (O) shown in Figures 2 to 8, as well as the concentrations of these elements shown in Table 1 for certain areas of the edge of the L fortunei shell, indicate that calcium phosphate occurs inside (Figures 7 and 8) and on the inner and outer faces of the pallial surface of the L fortunei mantle. We did not observe any overlap between carbon and calcium either inside or on the outer surface of the L fortunei mantle. Calcium carbonate (CaCO_3_) clearly appears in the nacreous layer (maps in Figures 7.a.3 and 7.b.3). In the images in Figures 7.a.1 to 7.a.4, calcium phosphate (phosphorus and calcium maps) forms a continuous film of ∼3 μm of thickness separating the epithelial face of the mantle from the nacreous layer. In 7.a.3, this film containing phosphorus and calcium is made up of particles which are in dissolution mode and self-organize below into the plates of the nacreous layer. The calcium map in 7.4.a shows the calcium in the CaCO_3_ layer and in the phosphate film.

Since the shell of the L fortunei is predominantly CaCO_3_, as shown by the results of ICP-OES, WDXRF, FTIR spectroscopy and X-ray diffraction (XRD), there must be processes within the calcium phosphate film that promote the replacement of phosphate into calcium carbonate, CaCO_3_, in both aragonite (nacre and innermost prismatic layer of the L. fortunei shell) and calcite phases (35).

Tears in the inner surface of the mantle of L fortunei shown in Figures 8.b.1, b.2, b.4 clarify that the inner surface of the mantle is ciliated, and that the interior of the mantle has a complex fiber structure. The carbon (C) map marks organic fibers and microscopic aggregates (one of them marked with a magenta asterisk), possibly of mucus, inside the mantle. Note that neither calcium nor phosphorus marks these forms (compare with the secretions coming from the cilia of the inner fold of the mantle in Figure 8.a.4). It is not possible to say that this mucus appeared inside the mantle because it could have been brought there during sample preparation (see Figure 1 for an overview of the areas examined).

The images of the edges confirm and describe in more detail the growth process of the L fortunei shell. Figure 2 shows images of the growth margin at the tip of the shell opposite the umbo. The damaged periostracum at this location made it possible to observe the biomineralization just below (Figure 2.a.1). Maps of Ca, C, P and O show that calcium and phosphorus mark the morphologies shown in Figures 2.a.2, a.3 (surface topography), 2.c.1 and c.3 (images with greater depth of focus). The striking similarity of these morphologies to the morphology of hydroxyapatite (HAp) can be seen in some studies in which HAp was obtained with a plate like morphology (56,57). The Ca/P ratio is used to characterize the type of calcium phosphate compound, and the value of 1.66 obtained by EDS quantification coincides with the value of the Ca/P ratio for HAp as shown in Table 1 (44,58).

The images in Figures 3.a and 3.b are magnifications of the plate morphology. The growth of the plates may be due to the adhesion of nanoparticles, which appear in large quantities in the images. The images in 3.c and 3.d were obtained from another L fortunei shell individual and confirm the presence of calcium phosphate containing composite plates at the growth edge (EDS maps in Figure 3.e). The thickness of the HAp layer in Figure 3.d is of the order of ∼18 μm and the thickness of the total calcium phosphate layer is of the order of ∼120 μm. Note in the detail of Figure 3.d that a surface is formed on the first HAp layer (marked with the red asterisk). This type of structure should facilitate mechanical stiffness of the growing tip.

One question that remains is whether this layer of calcium phosphate, whether it is HAp or carbonated hydroxyapatite (CHAp) or some other type of phosphate, will remain stable or whether it will be replaced by CaCO_3_ as the shell grows. XRD did not indicate the presence of HAp. However, the main XRD peaks of HAp occur in the same region as the main peaks of aragonite and calcite, the only phases identified by XRD in the L fortunei and P perna samples.

The SEM images, maps, and EDS compositional graphs shown in Figure 4 are from the A2 region of the L fortunei shell (Fig. 1.b). This is a region farthest from the tip of the growing shell. Again, fractures in the periostracum made it possible to analyze the composition and morphology of the calcium phosphate aggregates that appear at the edge of the shell, just where the periostracum curves and joins the edge of the mantle from which it originated. The aggregates do not have the same morphology as seen in Figure 2.a.3. Details of this morphology are shown in Figure 5.a.1 and show that below the calcium phosphate aggregates, a layer of plates (5.a.2 and a.3) is formed that is very similar to the CaCO_3_ nacreous layer of the L fortunei shell. The nanoparticles circled in red in the image of Figure 5.a.3 may be incorporated into the nacre to form this layer. The void that allowed the nacre layer to be seen could be the result of the dissolution of the micrometric particles shown in 5.b.1, which have a large surface area characterized by indentations and which could react with substances secreted by the surface of the mantle from which calcium phosphate was also secreted.

The presence of CA, or bivalve proteins with the same capability, in the mantle of mollusks and its role in biomineralization has long been discussed (36,39,59,60). Since the reversible reaction involving carbonic anhydrases is essential for many physiological processes, the presence of these zinc (Zn)-containing metalloenzymes is common throughout the animal, plant, and bacterial kingdoms. These enzymes accelerate the reversible hydration of carbon dioxide (CO_2_) to bicarbonate (HCO ^-^), a process that releases H^+^ and regulate the formation of CaCO_3_. CA acts in bone resorption by dissolving CHAp (carbonated hidroxyapatite) (61). A similar process may occur in the replacement of calcium phosphate particles by aragonitic CaCO_3_. The calcium phosphate film shown in Figure 7.a.4 may indicate a more advanced stage in the process of dissolution to form the nacre plates.

Miyashital et al(62) showed that nacrein inhibits the growth of calcium carbonate. In this work we show that enzymes dissolve calcium phosphate at the edge of the L fortunei shell and that it is from this dissolution/precipitation that calcium carbonate arises. The biomineralization of the shell of the bivalve L fortunei is a source of phosphorus compounds released into the animal’s organism as it biomineralizes and grows.

In L fortunei, the transformation at the edge of the shell (calcite), which causes the shell to grow in area, is not the same as that which occurs in the thickness of the shell (aragonite). At the edge there is a transformation of a morphology identical to the morphology of HAp. This morphology undergoes dissolution and continuous surfaces perpendicular to the plates grow, through the dissolution of the phosphate phase, to give way to the cubic morphology of calcite. A suggestion of how this process takes place is shown in figure S5 Supplementary Information.

The SEM images and compositional maps shown in Figure 6 are from the A3 region of the L fortunei shell (Figure 1.b). In this case, there are two fractures at the edge of the shell, just below the periostracum. In images 6.a.1 to 6.a.3 it is possible to see the presence of a layer containing phosphorus and calcium, but it was not possible to characterize what type of morphology was present. However, in images 6.b.1 to 6.b.4 it is possible to see that the morphology of the particles is the same as that seen in Figure 3. An analysis of these images indicates that there is a process of coalescence of plate morphologies. This coalescence of the plates may be due to the action of substances with dissociative capacity.

The images from 2 to 8 show that at the edges of the shell growth, phosphorus and calcium mark morphologies that appear to be in some process of transformation. The XRD and FTIR results from this study and previous ones (35,63) indicate that the shells of both L fortunei and P perna are composed of CaCO_3_ mostly in the aragonite phase (≥97%) and only the small remainder in the calcite phase. A micrometric film of calcium phosphate was identified at the interface between the mantle and the nacre of the face (Figure 7). Below this, the nacre appears to be built up from the aragonite phase (Figure 7.a.4 and EDS maps). The 3 μm thick layer (figures S6 in the supplementary material) consists of spherical particles of approximately nanometric size. At the edges, the presence of morphologies with striking similarity with crystalline hydroxyapatite in plate morphology(49) could be recorded in detail due to fractures in the periostracum. The molar ratio of Ca/P for plaque morphologies (refer to Table 1, lines 2 and 4) closely approximates the calculated value of 1.67 for hydroxyapatite and aligns closely with the proposed Ca/P (weight %) ratio for carbonated HAP (44).

The process of replacing any type of calcium phosphate by CaCO_3_ has not yet been reported in adult bivalves. However, several scientific studies from the first half of the 20th century confirmed the presence of calcium phosphate in the mantle and tried to understand the mechanism according to how calcium phosphate was replaced by CaCO_3_, as in the case of G Bevelander Bevelander(21,64). The presence of abundant calcium phosphate in the mantle, as discussed by authors in the early 20th century, was not consistent with descriptions of a mantle devoid of calcium phosphate (21). In 1938, Greenwald (65), quoted by Bevelander, intuited that the solubility of calcium phosphate increased in the presence of organic molecules. Interestingly, the enzyme carbonic anhydrase had been identified in 1933 (66), but its presence in the mantle of mussels was not yet known. In a 1952 article, Bevelander (22) used the radioactive isotope ^32^P to detect the presence of phosphorus in the shell and concluded that the phosphate must be in the periostracum. In this experiment, he also observed that the phosphorus came from the water (both fresh and marine) and that the phosphate occurred at the edges of the mantle near the mucus glands.

Our experiments with the shell of L. fortunei confirm the observation that calcium phosphate is located at the edges of the growing shell and, of course, next to the periostracum. To start the shell growth period, the mantle—a structure larger than the shell itself—must initially produce periostracum films (comprising at least three layers formed within the mantle groove(67,68) over the existing shell. These films need to firmly adhere to the existing periostracum before the biomineralization process can commence. In L. fortunei, the distances between the growth lines marked on the surface of the shells range from submillimeter to millimeters. In the case of the marine bivalve P perna, the growth lines are several millimeters apart because the shell is much larger than that of L fortunei. A diagram of the growth of the L fortunei shell is shown in Figure 9. The hydrophobic thin layers of periostracum are released from the outer fold of the mantle, having already polymerized, and adhere to the existing shell surface. In Figure 9, the dashed circle around the photo of the L fortunei shell shows the biomineralized saliency of this existing periostracum-shell overlap. XRD and FTIR results and previous work (35,63) indicate the presence of carbonate adhering to the periostracum. Substances secreted by the mantle, possibly enzymes (69,70), would then act to bring about the phosphate to carbonate substitution. On the right side of the diagram is the increase in the thickness of the mantle: the action of substances (marked with ★) transforms the nanoparticles that make up the calcium phosphate film shown in Figure 7 into aragonite plates.

What are the possible competitive advantages of the CaCO_3_ replacement process from the calcium phosphate precursor? The first, and perhaps most important, would be that the phosphate would be harvested mostly directly from the water, accumulated at the edges of the mantle (22), and then transferred to the pallial surface of the mantle in the form of nanoparticles or microparticles. This transformation would provide a source of phosphoric acids (61), which are essential for energy production (ATP), cell repair, pH regulation, and electric impulse transport. The mechanism of bone resorption may have its genetic ancestor in this stage of mantle construction(61).

Another advantage of the presence of crystalline calcium phosphate phases in the tip of the shell (images in Figures 2, 3 and 6) would be to ensure greater rigidity of the system in the process of biomineralization. The presence of amorphous phases would not provide the compressive strength that HAp possesses (71) and which allows the rigidity necessary for biomineralization to continue. The compressive strength of calcite and aragonite is lower than that of HAp.

The presence of a thin layer of crystalline HAp on the inner cuticle of bird eggs has been known for decades (72). The morphology of the HAp crystals also resembles the morphology found here in the periostracum of L fortunei.

In birds’ eggs, the HAp layer is the last layer (from the inside to the outside of the shell) and would paralyze the growth of calcite. However, subsequent work (73) has shown that there is phosphorus (in very low concentrations) throughout the thickness of the eggshell. Wouldn’t this phosphorus be evidence of a phosphate to carbonate replacement process that has taken place and that the thin layer of HAp on the egg cuticle indicates the end of the biomineralization cycle?

In L fortunei the calcium phosphate layer is the first to be deposited, after excreted by the mantle. Far from blocking the growth of CaCO_3_, the phosphate is dissolved and carbonate precipitates. This substitution would take place because the bivalve secretes certain substances like carbonic anhydrase and related enzymes that favors it. In the case of L fortunei, crystalline HAp is replaced by crystalline carbonate forms, either calcite or aragonite, at the tip of the shell (growth in length). When growing in thickness (nacre), the phosphate film must be amorphous and transform into amorphous CaCO_3_ nanoparticles (ACC) which, when aggregated, transform into the aragonite phase (metastable) in tablets.

Our study did not evaluate the presence of organic content (chitosan fibers, collagen or proteins). Such components may sum up to ∼5% of the composition of the shell. But their production and presence in the composite that makes up the shell must certainly be associated with the release of phosphate that have become energy sources to produce these composites in the mantle and elsewhere in the body of the bivalve.

**Figure 9.**
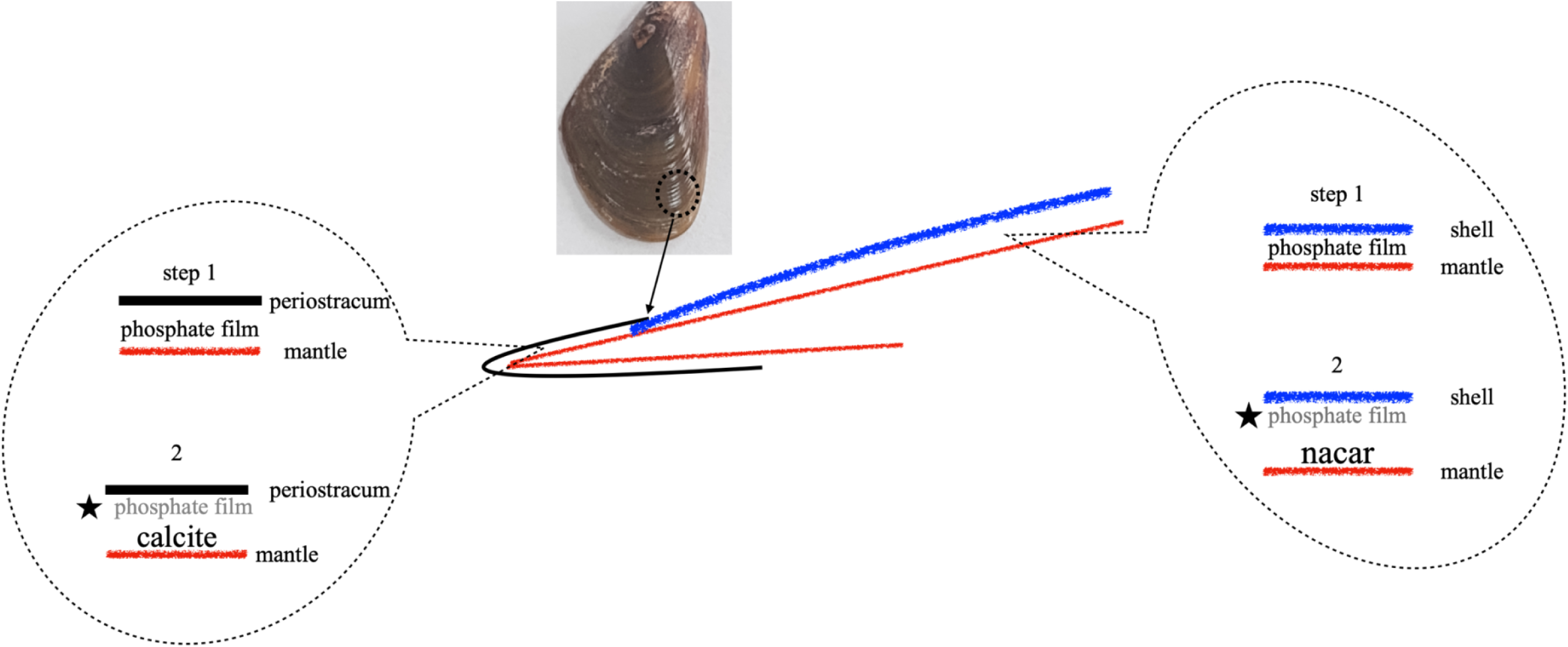
– Diagram of L. fortunei shell growth. Periostracum sheets (black line) are launched from the outer mantle fold (red line) and adhere to the existing shell surface (blue line). Distances are millimeters or submillimeter in the case of L fortunei. The dashed circle in the shell photo shows the biomineralized protrusion of this existing periostracum-shell overlap. On the left side of the diagram, the shell growing takes place: XRD and FTIR results of this and previous works (35,63) indicate the presence of calcite adjacent to the periostracum. The periostracum connects to the mantle at the bottom of the shell (see edges in Figure 1). The star (★) followed by a grey “phosphate film” words represent the action of substances (carbonic anhydrase or other enzymes) secreted by the mantle, which would act to bring about the phosphate to carbonate replacement. On the right side of the diagram is the increase in the thickness of the mantle: the action of the organic substances (★) transforms the nanoparticles that replaces the calcium phosphate film shown in Figure 7 by aragonite plates.

## 5. Conclusions

The mantle-shell interface and the shell edges of the freshwater bivalve L. fortunei have been studied using different experimental techniques, and it has been shown that the mantle, although millimeters thick in the golden mussel, has a complex internal structure where physiological processes and physicochemical replacements take place that have not yet been elucidated. SEM-EDS spectroscopy images of the shell edge revealed the presence of calcium compounds with both carbon (C) and phosphorus (P). In the shell edge of L. fortunei, the calcium phosphate phases are the first to be deposited in the periostracum, both in the growth of the shell area and in the thickness of the shell through the formation of the nacreous layer. The process of substitution of phosphate into calcium carbonate must involve proteins that contain the carbonic anhydrase (CA) function, which provides protons for the dissolution of calcium phosphate and bicarbonate for the formation of calcium carbonate. In the case of L. fortunei, this process takes place on a micrometric scale, with the phosphate film on the epithelial side of the mantle being ≍ 3 μm thick.

In the growth of the edge of the shell tip (Figures 2 and 3), calcium phosphate compounds whose morphologies are very close to the hydroxyapatite (HAp) crystalline phase were identified as the first phase that mineralizes at the bottom of the periostracum. The plates of HAp or similar compounds tend to coalesce to form surfaces (red asterisk in figures 2.c.2, 3.d and 6.b.4). The presence of crystalline phases should help to stiffen the growing edge. The way the transformation into calcite occurs is suggested in figure S5 (Supplementary Material). This work shows that calcite forms a micrometric film adhering to the periostracum (figure S4, Supplementary Material).

In the L fortunei shell region that grows in thickness, the presence of a calcium phosphate film composed of nanoparticles was observed (Figures 7.a.3, 7.a.4 and S6 in Supplementary Material). Immediately below this film is the nacreous layer with tiles (or bricks) characteristic of the aragonite morphology. There is a clear separation between the epithelial surface of the mantle, the calcium phosphate nanoparticle film, and the nacre. Images at the edge of the lateral part of the shell (Figures 4 and 5) show aggregates labeled as calcium phosphate next to the periostracum and, just below, the formation of tiles (or bricks) labeled with calcium (Ca) but no longer with phosphorus (Figure 4.b.1).

The composition of the shells of the dulcicolous bivalve L fortunei and the marine P perna by ICP-OES and WDXRF indicate the concentration in ppm for the phosphorus (P) present in the shell, showing that the presence of calcium phosphate phases in the shell is limited to micrometric films in the process of being replaced by calcium carbonate. ICP-OES tests on the shell carried out on different occasions show virtually identical concentrations regardless of the year of collection.

For XRD, FTIR and FTIR-ATR spectroscopy, pieces of the shell edges of the two species of bivalve mollusks were pulverized. In the case of FTIR-ATR, whole pieces of the shell edge of the two mussels were analyzed. The absorption band of the phosphate group around 1020 cm^-1^ was tentatively assigned in both FTIR and FTIR-ATR spectroscopies. Carbonate however is the phase with clearly defined absorption bands.

XRD confirmed the presence of aragonite and calcite in the shells. Aragonite occurs in high concentrations (over 90% of the crystalline phase) and calcite in much lower concentrations in both bivalves. Due to the concentration, no calcium phosphate phases were detected. However, one of the main peaks of the calcium phosphate crystalline phase, 2θ=25.86° (002), is present in the L fortunei (2θ=26.26°) and P perna (2θ=26.21°) samples.

The presence of phosphate as the primary phase in the biomineralization process of L fortunei reopens the discussion on the importance of the phosphate to carbonate replacement during the biomineralization of bivalves and indicates an important evolutionary advantage by releasing phosphate compounds essential for energy production and the other metabolic functions of the organism. The presence of calcium phosphate aggregates inside the mantle was detected by EDS spectroscopy, but it was not possible to confirm whether these aggregates were absorbed by the inner ciliated surface of the mantle. The connections between the mantle and the circulatory system of L. fortunei could, in future studies, verify the transit of phosphate in the organism. In these future studies, the SEM-EDS study of the marine bivalve P perna could confirm the importance of the phosphate phases in its biomineralization. XRD, FTIR and FTIR-ATR spectroscopies and chemical analysis by ICP-OES (quantitative) and WDXRF (semi-quantitative) indicate the presence of phosphorous and phosphate in the shell of P perna.

## ACKNOWLEDGEMENTS

To the Microscopy Center of UFMG (Federal University of Minas Gerais) and to the Crystallography Laboratory of the Physics Department (LabCri) of UFMG for their support in carrying out this study. A special thanks to Professor Leonardo H. R. Dos Santos, Crystallography Laboratory of the Chemistry Department of UFMG, for his help and support both for reading this work and helping on the analyses of the FTIR and ATR experiments and results.

