## Supplemental Figs S1 to S7 for "Dissolution of Phosphate and Precipitation of Carbonate in the Biomineralization of the Bivalve Shell Limnoperna fortunei"

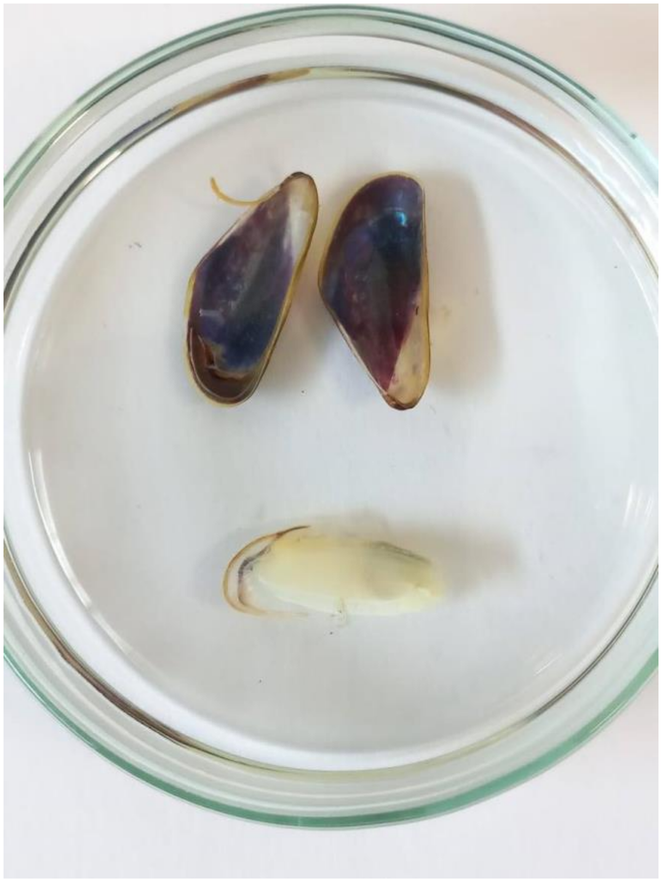

**Figure S.1**- The upper part of the photo shows the two shells of the bivalve Limnoperna fortunei. The lower part of the photo shows a mantle (pallium in Latin) of the golden mussel. Each shell arises from the biomineralization produced by the mantle of the bivalve.

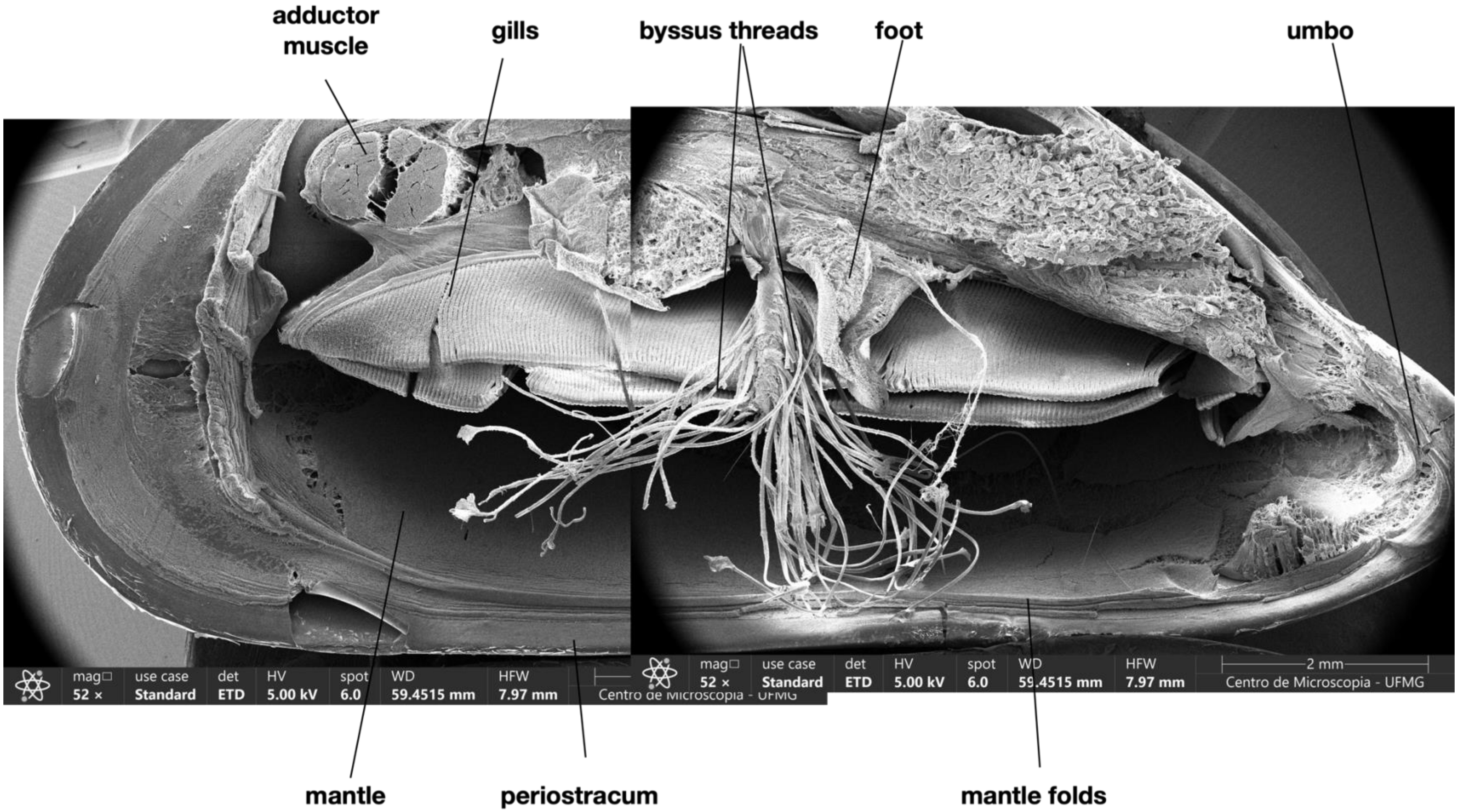


**Figure S.2 -** Anatomy of L. fortunei, showing the parts that are relevant to this work.

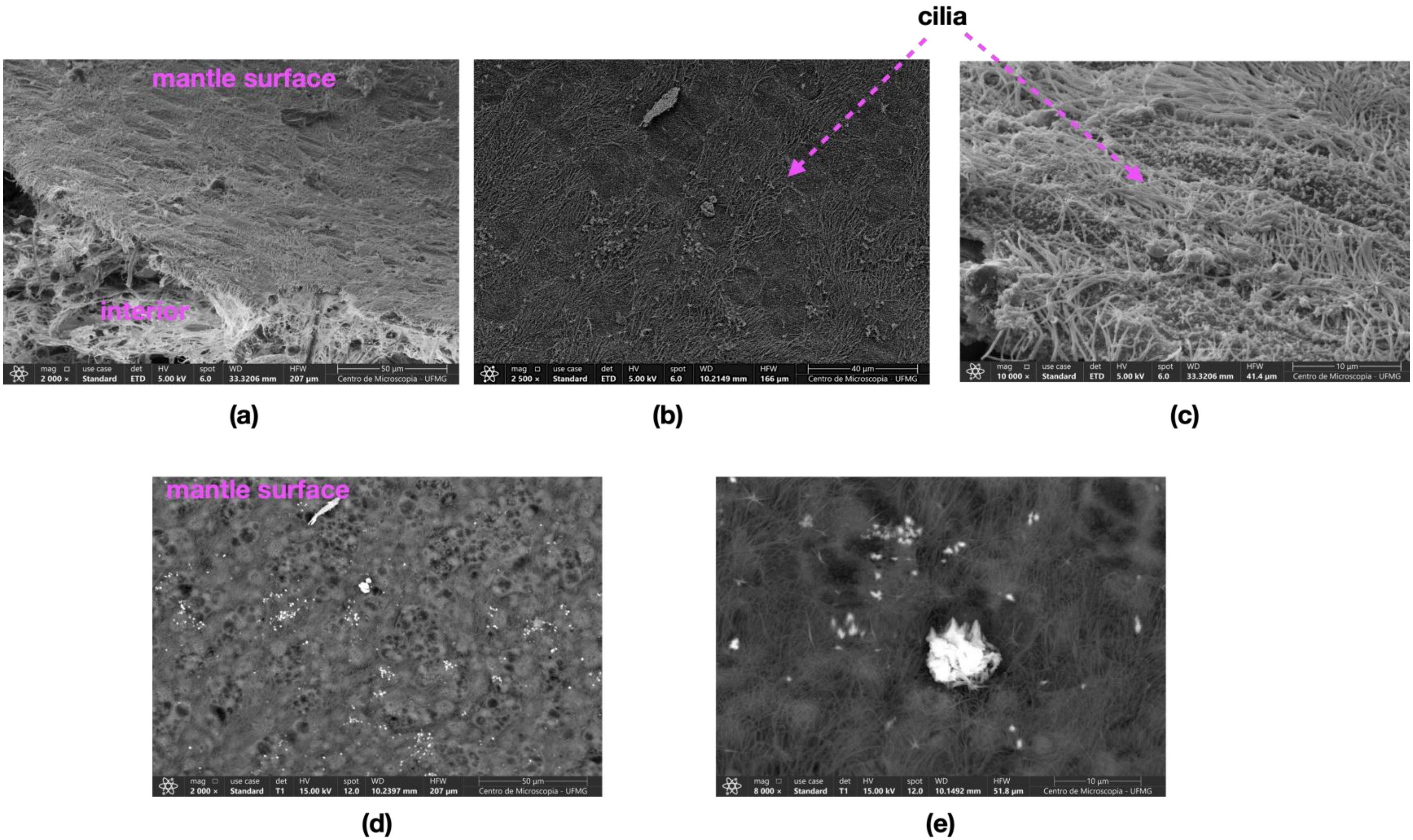


**Figure S3 – Internal** surface (facing the gills) of the L. fortunei mantle at different magnifications: (a) Displays the internal surface and interior of the L. fortunei mantle; (b) Cilia patches are evident on the internal surface of the mantle; (c) Detail of the cilia, with some areas devoid of cilia visible; (d) Using a SEM-T1 detector, black circles that appear to be micrometer-sized pores are observed on the mantle's surface; (e) Detailed view of these pores with a particle situated on one of them. Further studies will investigate the nature of these openings to determine whether they are artifacts from sample preparation or genuine pores in the mantle wall.

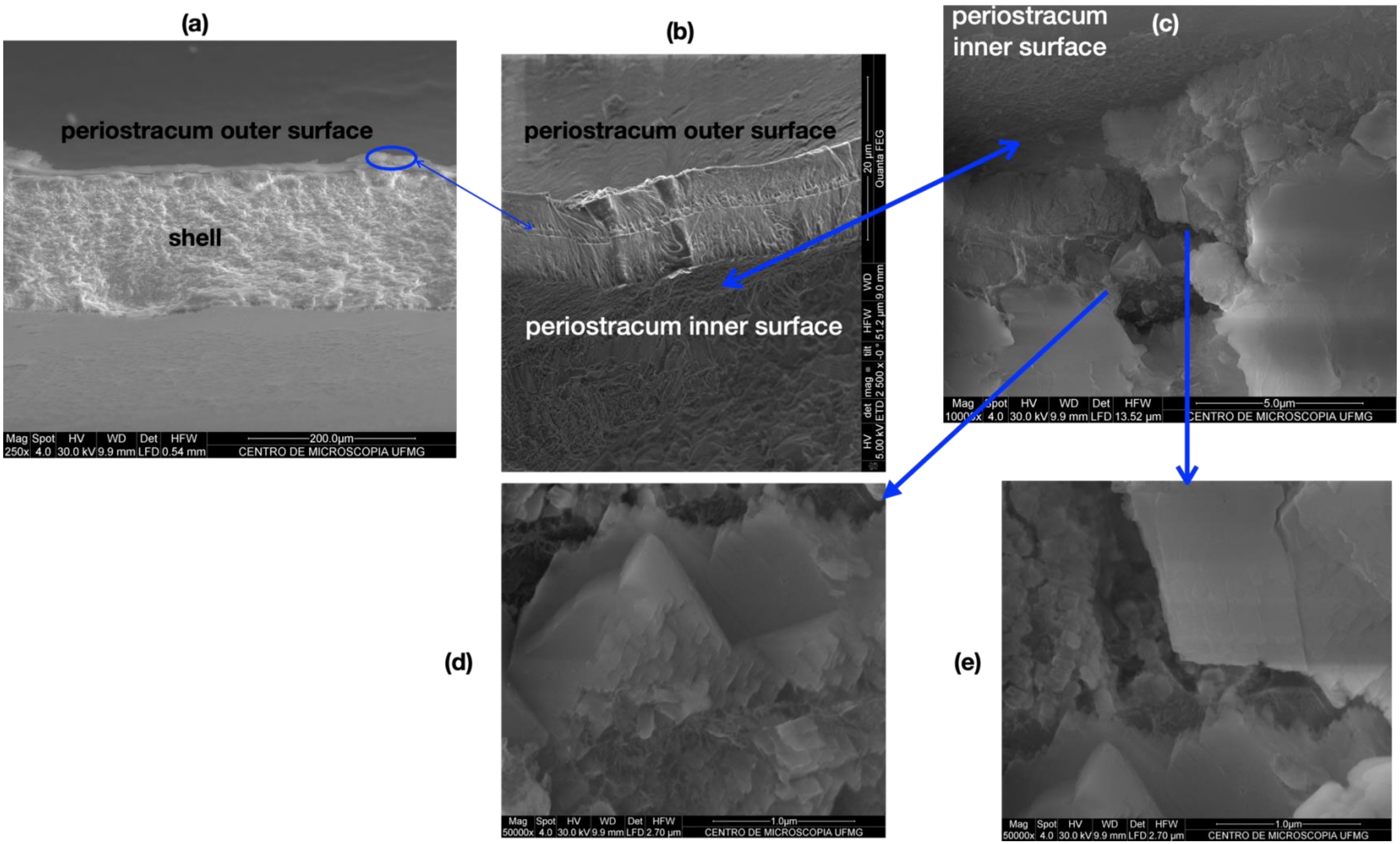


**Figure S4** – (a) Broken L. fortunei shell displaying the entire cross-section of the shell. Periostracum circled in blue; (b) Periostracum of L. fortunei featuring its two layers and the calcite layer beneath it; (c) Detail of the calcite layer below the periostracum; (d) and (e) Detailed views of the microstructure of the calcite layer.

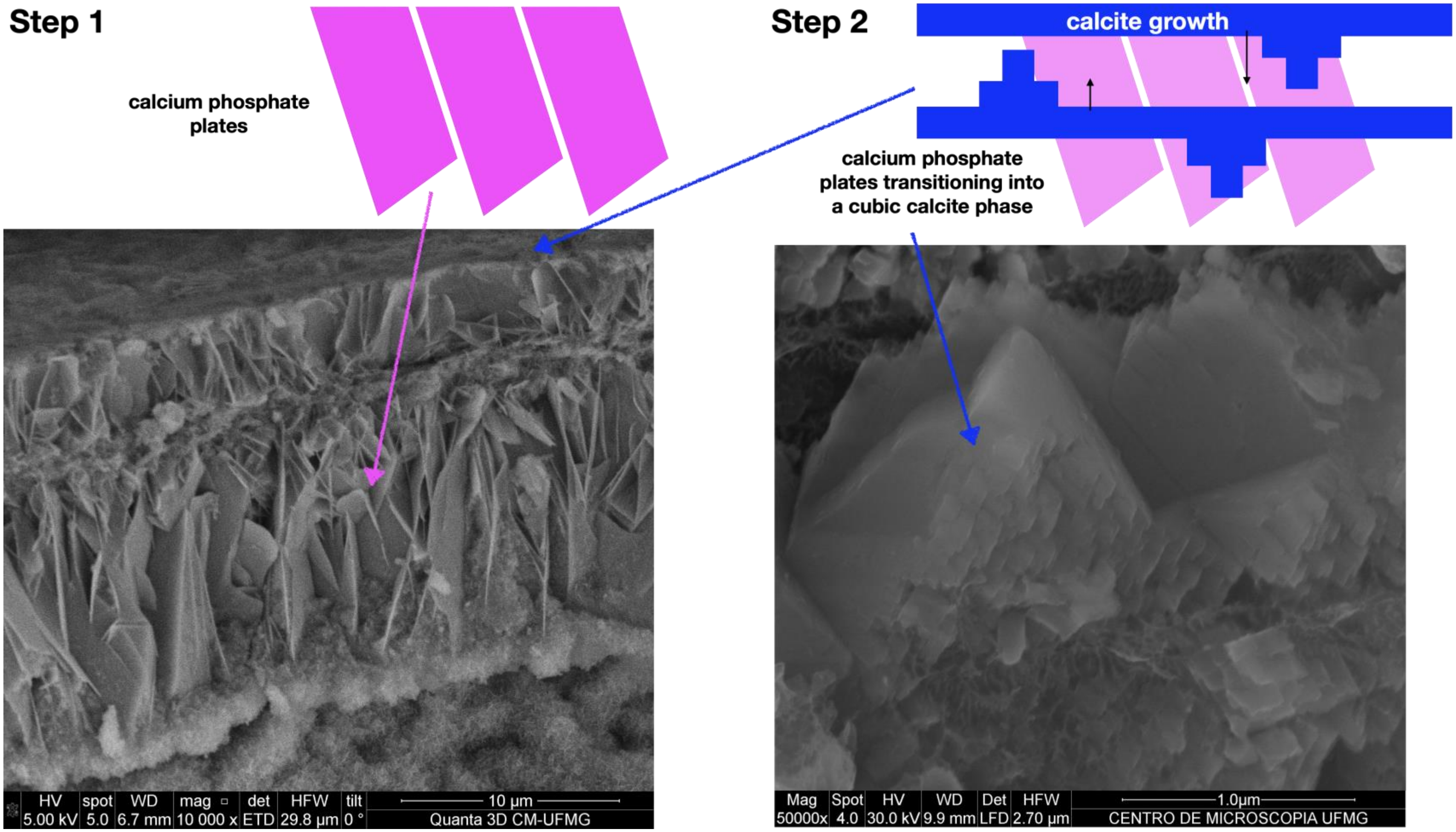


(a) (b)

**Figure S5-** Transformation of calcium phosphate -> calcite at the growth edge of the L. fortunei shell via enzimatic : (a) Calcium phosphate plates with morphology similar to hydroxyapatite (HAp) or carbonated HAp (CHA), dahllite (1) , exhibit coalescence, possibly through dissolution of calcium phosphate and precipitation of calcium carbonate. (b) Monocrystals or polycrystals of calcite present immediately below the periostracum of the L. fortunei shell. See location in the images of Figure S. Above each image, diagrams appear (step 1 and step 2) suggesting the process of dissolution of calcium phosphate and precipitation of calcium carbonate in the calcite phase. Dissolution takes place by protonation of the carbonate group to form carbonic acid and protonation of the phosphate groups to form phosphoric acid (2).

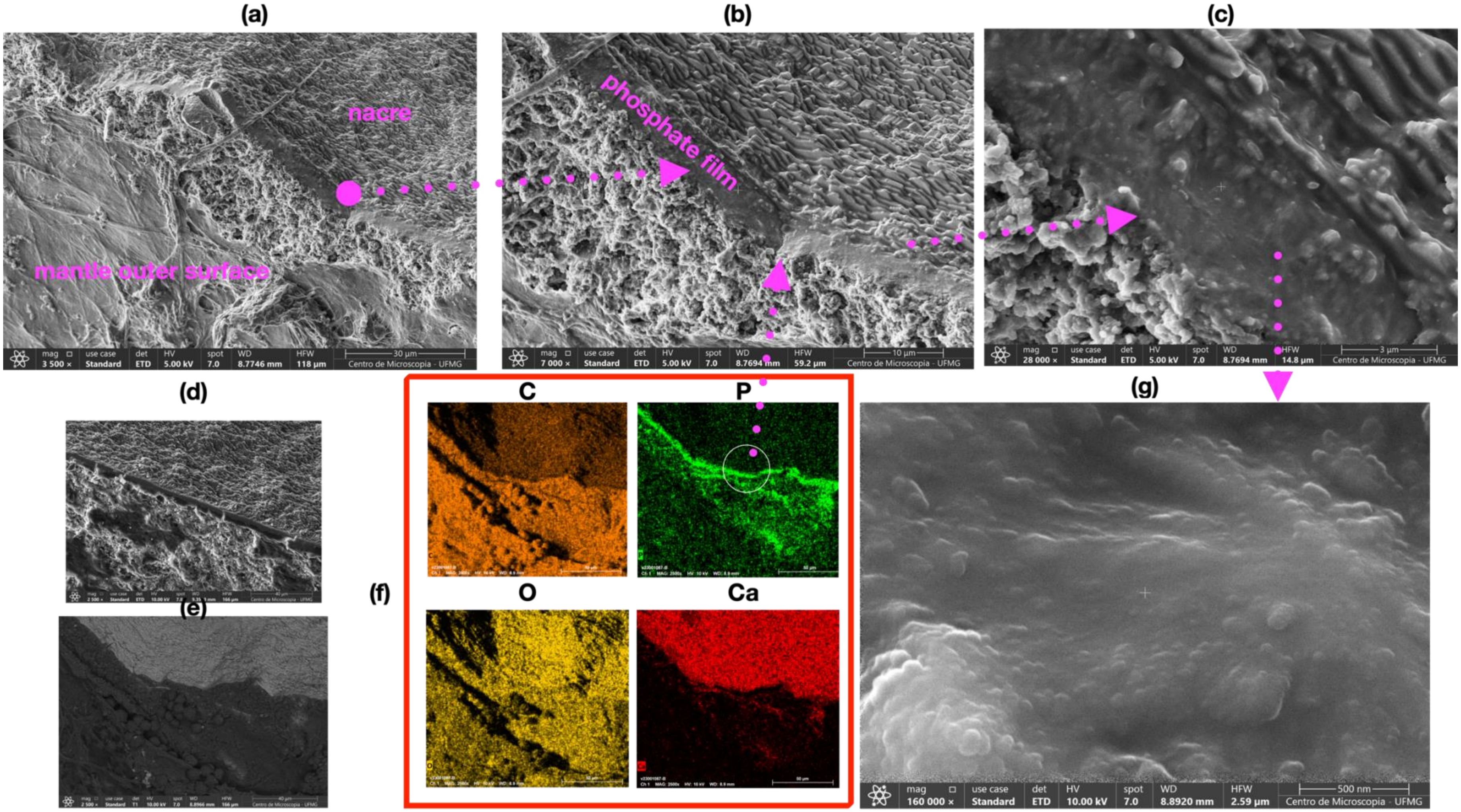


**Figure S6-** (a) Surface of the mantle (lower part of the image) and nacre in the upper part of the image; (b) Between the mantle and the nacre the phosphate film; (c) Image of the phosphate film in detail. (d) and (e) show the same region using different SEM sensors; (f) Maps of the four main chemical elements observed in the image: carbon (C), phosphorus (P), oxygen (O) and calcium (Ca); (g) Nanoparticles in the phosphate film.

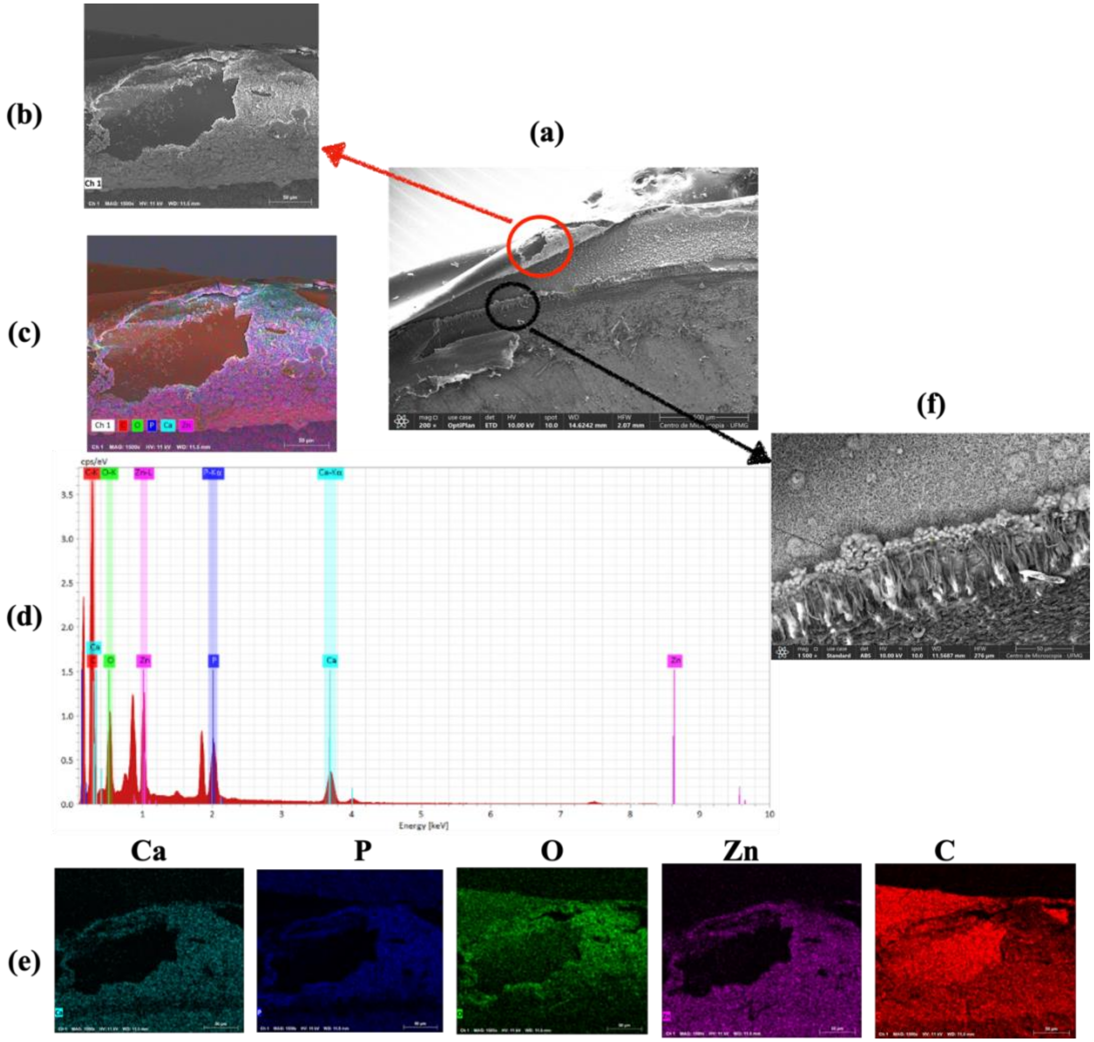


**Figure S7** - (a) Sample from the edge of the L fortunei shell (SEM) was analyzed by EDS; (b) Area circled in red is shown in detail. (c) The main chemical elements found were C, Ca, P, O and Zn, as shown in the atomic concentration graph in (d). (e) The element maps show that Zn marks the region where calcium (Ca), phosphorus (P) and oxygen (O) occur. Carbon (C) does not coincide with the other elements. (f) Shows in detail the region circled in black. The morphology of the crystals in the upper part of this image is identical to that shown in figures 2, 3 and 6 of the article. The presence of zinc (Zn), a metallic element in the enzyme carbonic anhydrase (CA), was not found in other samples, only at this edge.
